# A proteome-wide structural systems approach reveals insights into protein families of all human herpesviruses

**DOI:** 10.1101/2024.07.16.603793

**Authors:** Timothy K. Soh, Sofia Ognibene, Saskia Sanders, Robin Schäper, Benedikt B. Kaufer, Jens B. Bosse

## Abstract

Structure predictions have become invaluable tools, but viral proteins are absent from the EMBL/DeepMind AlphaFold database. Here, we provide proteome-wide structure predictions for all nine human herpesviruses and analyze them in depth with explicit scoring thresholds. By clustering these predictions into structural similarity groups, we identified new families, such as the HCMV UL112-113 cluster, which is conserved in alpha-and betaherpesviruses. A domain-level search found protein families consisting of subgroups with varying numbers of duplicated folds. Using large-scale structural similarity searches, we identified viral proteins with cellular folds, such as the HSV-1 US2 cluster possessing dihydrofolate reductase folds and the EBV BMRF2 cluster that might have emerged from cellular equilibrative nucleoside transporters. Our HerpesFolds database is available at https://www.herpesfolds.org/herpesfolds and displays all models and clusters through an interactive web interface. Here, we show that system-wide structure predictions can reveal homology between viral species and identify potential protein functions.

## Introduction

The *Orthoherpesviridae* family comprises alpha-, beta-, and gammaherpesviruses containing nine human herpesviruses (see Table 1 for virus abbreviations and their respective subfamilies). These are medically important human pathogens, including viruses such as human cytomegalovirus (HCMV), the leading infectious cause of congenital disorders in the USA. Due to the large number of encoded genes (between 70 and 181, depending on the species), experimental structural information for most viral proteins is not available, severely hampering their functional analysis. Moreover, herpesvirus gene and protein nomenclature are very inconsistent between species. Even worse, some proteins of the same species have multiple names based on their genome position, molecular weight, expression in the infected cell, or packaging into the virion. This situation makes it difficult even for experts to draw parallels between viruses. While some corresponding orthologues between species can be deduced from the UniProtKB annotation, this database is incomplete and sometimes inconsistent. Finally, herpesviruses also have a penchant for duplicating genes, further complicating annotation.

**Table 1.** Viral proteins used for structure prediction.

| Subfamily | Virus | Virus abbreviation | Strain | Proteome ID or Reference Sequence | Number of proteins |
| --- | --- | --- | --- | --- | --- |
| alpha | herpes simplex virus 1 | HSV-1 | 17 | UP000009294 | 77 |
| alpha | herpes simplex virus 2 | HSV-2 | HG52 | UP000001874 | 74 |
| alpha | varicella-zoster virus | VZV | Dumas | UP000002602 | 70 |
| beta | human cytomegalovirus | HCMV | Merlin | UP000000938 | 181 |
| beta | human herpesvirus 6A | HHV-6A | U1102 | NC_001664 | 90 |
| beta | human herpesvirus 6B | HHV-6B | Z29 | NC_000898 | 97 |
| beta | human herpesvirus 7 | HHV-7 | RK | NC_001716 | 84 |
| gamma | Epstein-Barr virus | EBV | B95-8 | UP000153037 | 87 |
| gamma | Kaposi's sarcoma-associated herpesvirus | KSHV | GK18 | UP000000942 | 86 |

With the development of machine learning-based prediction software^1,2^, the accuracy of the structural information has increased so that it can often be directly used for building wet-lab testable hypotheses, making it an invaluable tool for molecular biologists^3^. DeepMind, in cooperation with EMBL-EBI, has published over 200 million protein structures to facilitate this process, covering most of UniProtKB in an open-access database (https://alphafold.ebi.ac.uk). While this is an extraordinary resource for many fields, it currently excludes viral proteins such as the human herpesvirus proteomes.

Herpesvirus phylogeny has classically been sequence-based, such as a recent study that classified the human herpesvirus proteomes into homology families^4^. However, the current revolution in structure prediction and rapid structural similarity search algorithms^5^ has vastly expanded the pool of sequences that can be interrogated by structure-based phylogeny^6–8^. When applied to large proteins or large phylogenetic distances, *e.g.* across the *Duplodnaviria* realm, this approach clustered species more consistently than the sequence-based ones^9^. Moreover, a recent study using this approach showed that orthopoxviruses acquired cellular proteins, with some losing their original function^10^. Furthermore, structure prediction of the glycoproteins of different flaviviruses identified novel and common mechanisms in the fusion machinery^8^.

Here, we predicted the structures of all nine human herpesvirus proteomes, totaling 844 individual proteins. We critically evaluated their accuracy with stringent quality scores and thresholds. We clustered their predicted structures into structurally similar groups and analyzed them using a structural systems virology approach. Many similarities suggest a common ancestor and thus a homologous relationship, whether orthologous, *i.e.* due to speciation, or paralogous, *i.e.* due to gene duplication. We found new members to previous groups identified by sequence, such as the US22 family of HCMV. We could also identify unexpected groups with distinct folds, such as the UL112-113 family harboring a unique beta-barrel domain that is conserved in seven of the nine human herpesviruses. A detailed domain-level similarity search identified several subclusters of proteins harboring different numbers of duplicated domains, deletions, and acquisitions. Using structural similarity searches against cellular proteins, we propose new functions for uncharacterized proteins. Examples include the HSV-1 and HSV-2 US2 proteins, which potentially have dihydrofolate reductase activity. We also found cases of exaptation, such as the BMRF2 family of transmembrane proteins that share structural similarity to nucleoside transporters but are likely inactive and now have functions in cell-cell spread.

In this work, we create a structural database that is accessible through an open-access and searchable web interface: https://www.herpesfolds.org/herpesfolds. It offers interactive displays of the predicted structures and structural clusters. HerpesFolds is a curated database that groups the protein predictions based on structural similarity while linking their established names and groups. It also serves as a reference that can translate the complicated herpesvirus nomenclature for the expert and non-expert alike and sheds light on the relationships of all nine human herpesvirus proteomes. This work will aid in developing wet-lab testable hypotheses and gain new insights into herpesvirus biology and pathology.

## Results

### Most herpesvirus proteins can be confidently predicted

To generate proteome-wide predictions of prototypic strains of all 9 human herpesviruses (Table 1), we initially used LocalColabFold^11^, an implementation of AlphaFold2 that mainly uses a 40-60x faster multiple sequence alignments (MSA) generator (Figure 1A). For each protein, 5 models were generated. For most proteins, the associated predicted local distance difference test (pLDDT) plots of all five models were very similar. This indicated that the models converged in all five runs to a similar solution, indicating that the best solution was found (Supplementary Fig. 1A). To categorize the models objectively, we established three thresholds based on the pLDDT consistency between models and the predicted template modeling (pTM) scores. See the methods section for more details. Of the 844 proteins, 249 (30%) failed these quality scores (Figure 1B and Supplementary Data 1 Sheet 1). We used standard AlphaFold^1^ (Supplementary Data 1 Sheet 2) to improve these predictions using the slower but more thorough jackhmmer MSA generator algorithm as well as LocalColabFold^11^ with additional recycles (Supplementary Data 1 Sheet 3). This resulted in good predictions for 95 (38%) of the previously rejected models. Therefore, the final dataset consisted of 690 proteins (82% of 844 proteins) with consistent models as indicated by the pLDDT plot (Supplementary Data 1 Sheet 4). We compared the proteins that failed and passed the quality scores for each threshold criterion (Figure 1C-E). Most proteins that did not pass had low pTM and a small percentage of sequence with a pLDDT >0.70. This correlation suggested that proteins low in folded regions, as indicated by a pLDDT >0.70, had lower prediction confidence. The pLDDT score can also predict whether a protein region is structured or intrinsically disordered. As shown before, a pLDDT >0.70 indicates a well-modeled region with a Cα root mean square displacement likely less than 1.5 Å^12^. Conversely, pLDDT scores <0.50 likely indicate intrinsically disordered regions (IDR)^1,13,14^. Consequently, the AlphaFold database has been used to predict protein disorder, such as in MobiDB^15^. IDRs have recently been implicated in essential processes such as replication compartment formation of HCMV by liquid-liquid phase separation^16^. Therefore, we determined which well-predicted proteins likely contain disordered regions by quantifying the percentage of pLDDT values each model has <0.50 or >0.70. As shown in Supplementary Fig. 2A, 197 (23%) of the predicted proteins likely contain a disordered region (cut-off >44% of total sequence), and 640 (76%) of the proteins are likely structured (cut-off >33% of total sequence) (Supplementary Data 1 Sheet 5). This can be compared to the 1-8% disordered proteins in bacterial proteomes, 2-11% in archaea, 23-28% in eukaryotes, and 45% in humans^15,17,18^. Examples of N-terminal or C-terminal structured domains in a disordered protein are illustrated (Supplementary Fig. 2B). We also found a subset of 100 proteins that did not pass the pTM threshold but showed converging model pLDDT plots, indicating that the predictions came to a final solution. These proteins contained extensive sequence stretches with a pLDDT <0.50, which indicated that these proteins are well-predicted, disordered proteins. Finally, to indicate transmembrane domains as well as signal peptides, all predictions were further annotated with DeepTMHMM (Supplementary Data 1 Sheet 6).

**Figure 1.**
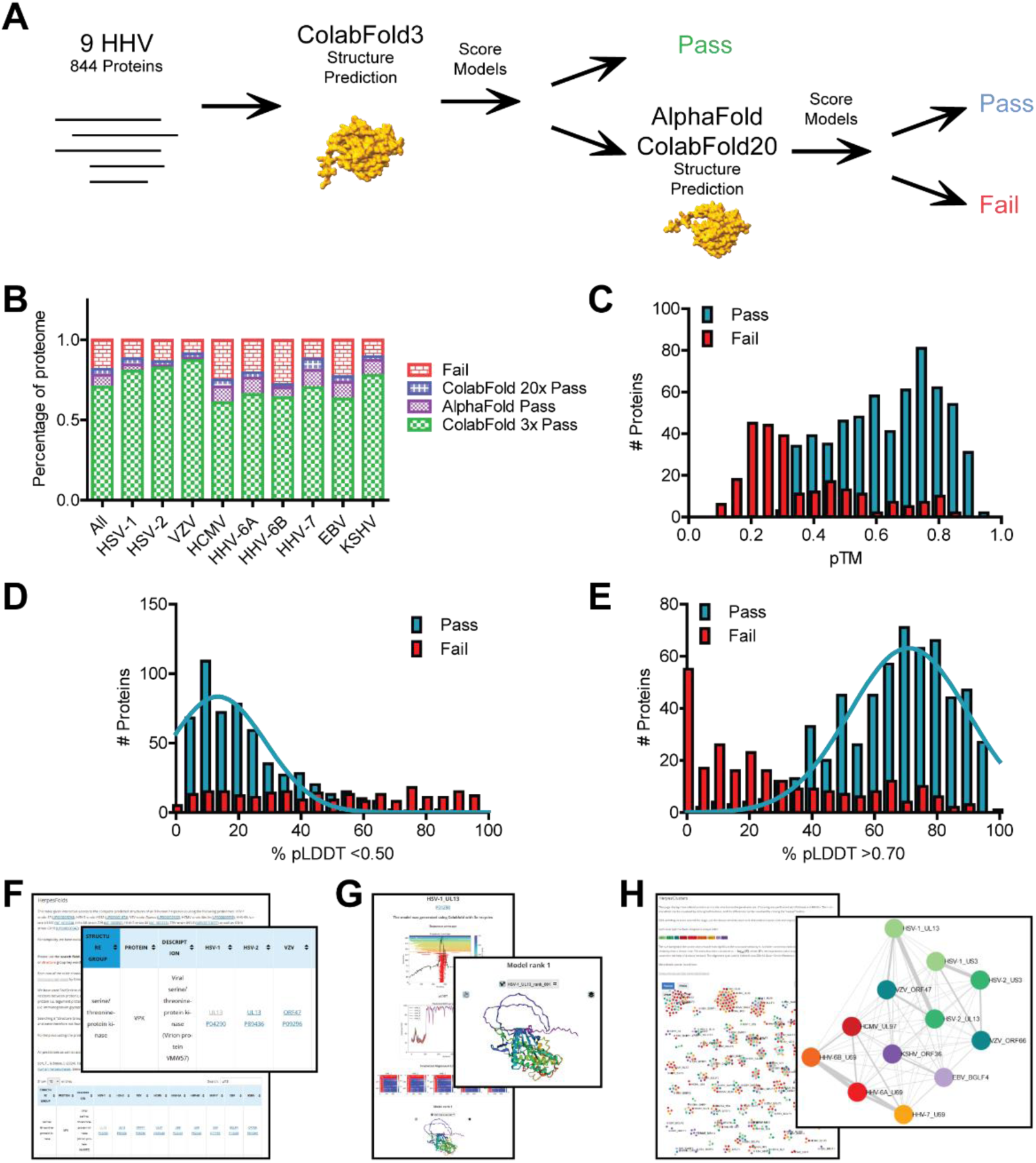
Structure predictions of the proteome of all human herpesviruses. A. Pipeline of model generation and quality scoring. The structures of 844 proteins covering the proteomes of all nine human herpesviruses were initially predicted with LocalColabFold with 3 recycles. After scoring, models that failed any quality score were rerun with AlphaFold and LocalColabFold with 20 recycles. B. Percentage of each viral proteome that passed the quality score. C. Histogram of the pTM scores of the initial LocalColabFold models that passed or failed the quality scores. D-E. Histogram of the percentage of each protein with a pLDDT score below 0.50 (D) or above 0.70 (E). In both cases, a Gaussian distribution was fit to identify the distribution for the proteins that passed the quality scores. F. HerpesFolds online database available at https://www.herpesfolds.org/herpesfolds. Structure predictions are organized and searchable by homology and structural similarity. G. Example webpage of the displayed information for each predicted model. H. Searchable clustering of all human herpesvirus proteins based on structure (Foldseek) or sequence (HHblits) similarity is available at https://www.herpesfolds.org/herpesclusters.

To illustrate the accuracy of the general prediction workflow, we compared our predictions to experimental data that were released after the AlphaFold 2.3 training cut-off of 2021-09-30. Our predictions aligned very well with experimental structures, as depicted for HCMV UL77^19^ (Supplementary Fig. 3). The Cα RMSD (310 atoms) between the predicted and experimental HCMV UL77 structure (7NXP^19^) was 0.509 Å, indicating a highly consistent model. These observations are in line with a previous community assessment that concluded that AlphaFold models are, on average, as good as experimental structures^3^.

### HerpesFolds is a searchable online structure database

To make our proteome-wide predictions accessible and easily searchable, we created a database at https://www.herpesfolds.org/herpesfolds (Figure 1F). The main table contains the individual herpesvirus protein names and their UniProtKB accession numbers. Based on the previous knowledge^20,21^, we generated a chart where each row contains proteins with known homology (Supplementary Data 2 Sheet 1). This way, the database also serves as a reference guide. This feature is important as the nomenclature of human herpesviruses is highly inconsistent and difficult to interpret even for experts. We also integrated the common names based on the UniProtKB annotation. Moreover, each protein entry links to the UniProtKB webpage and an individual webpage containing an animated model and the pLDDT and PAE graphs (Figure 1G). If predictions were rerun, the best model is displayed, and if the models of all runs failed the thresholds, a warning is displayed at the top of the page. Furthermore, transmembrane domains as detected by DeepTMHMM are indicated and a second model lacking the signal peptide is provided. As an important feature, we also integrated a Foldseek^5^ search link kindly provided by the Steinegger lab that directly submits the predicted model to the Foldseek server and displays structurally similar proteins. Finally, we interactively display the structural similarity network (Figure 1H).

### Clustering of known and unexpected structural homologs

The identification of herpesvirus orthologs has classically been through sequence alignment, but this approach is restricted by its need for some extent of conserved sequence similarity. To find more distant relationships that might represent homologs between different viral species and determine the potential function of orphan genes, we employed structural similarity searches. We used both DALI^22^ as an established algorithm and the newer Foldseek^5^ algorithm and ran all models against each other. We used a threshold of DALI Z-score of >8 and E-value <0.001 for Foldseek (see materials and methods) (Supplementary Data 2 Sheet 4 and 5, respectively). Overall, the resulting structural clusters were similar, with 615 proteins in protein similarity clusters for DALI and 632 for Foldseek. One example of a difference is the tegument protein 3 (TEG3) group. Foldseek grouped only the alphaherpesvirus TEG3 proteins with the gammaherpesvirus BGLF3.5/ORF35 proteins. In contrast, DALI also found a similarity of the roseolovirus (HHV-6A, HHV-6B, and HHV-7) U68 proteins with this group. Since Foldseek performed slightly better overall and is computationally much more efficient, we used it for the remainder of this work. The resulting structural similarity connections were used to cluster the herpesvirus proteins into groups of structural similarity (Figure 2A). All clusters are available for interactive exploration at https://www.herpesfolds.org/herpesclusters.

**Figure 2.**
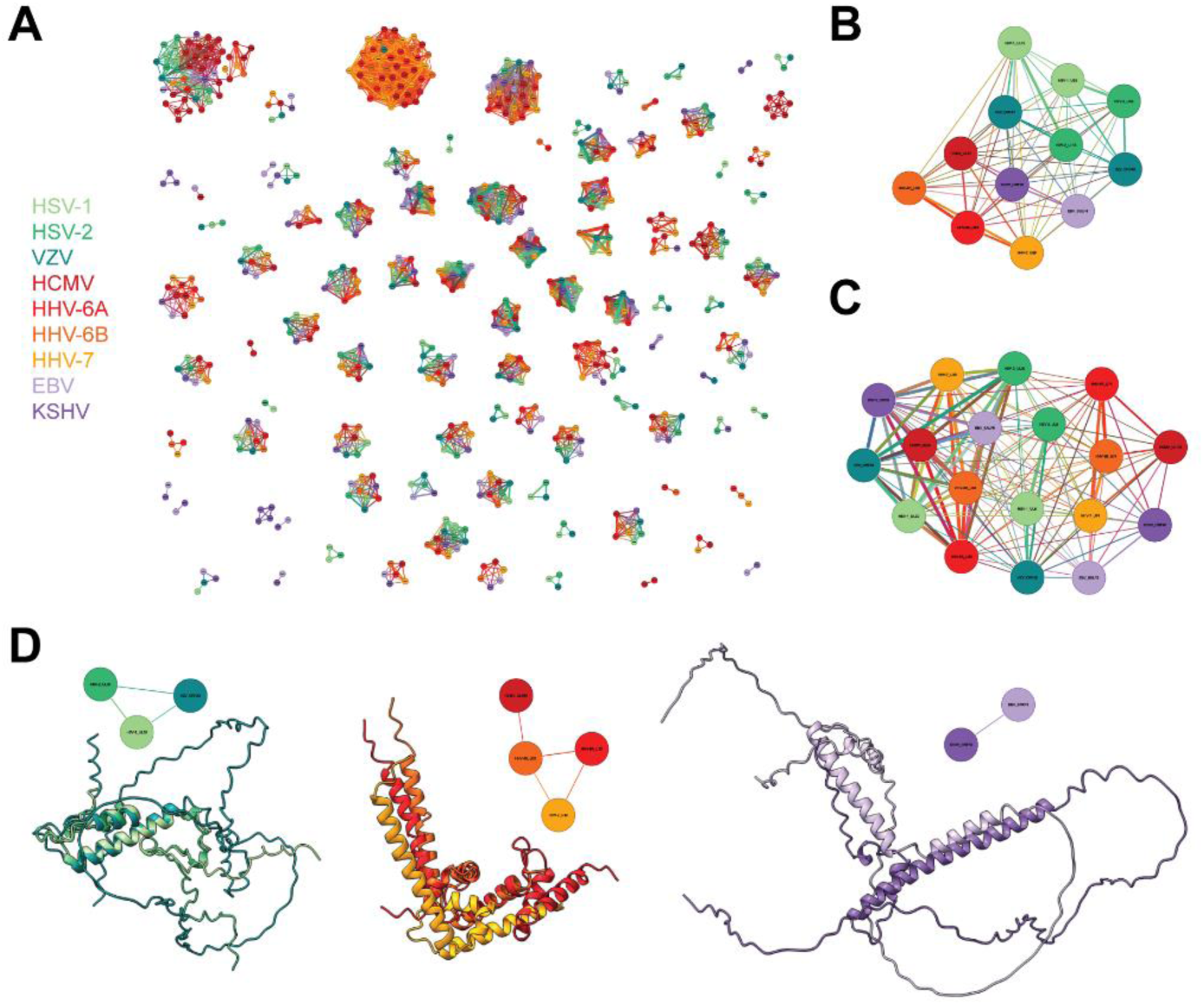
Structural similarity search identifies protein clusters. A. Graphical display of all structurally similar protein clusters. The 690 herpes proteins that passed the quality score were analyzed pairwise by Foldseek for structural similarity. The color scheme for the viral species is used throughout this work. The name of the protein is written in the center of each node and is legible upon zooming in https://www.herpesfolds.org/herpesclusters allows interactive browsing of the clusters. B. Cluster of viral serine/threonine-protein kinases US3 and UL13. C. Cluster of viral DNA helicase/primase complex-associated protein (HEPA) and DNA polymerase catalytic subunit (DPOL). D. Clusters of small capsomere-interacting protein (SCP). The alpha-, beta-, and gammaherpesviruses clustered separately. The corresponding UCSF ChimeraX session can be found at https://zenodo.org/records/13284140.

Based on the UniProtKB^20^ and previous work^4,21^, 39 protein groups were expected to be found in all 9 species. A more recent sequence analysis based on domain architecture identified 23 proteins with a consistent domain structure in all 9 species and 169 homology groups^4^. Using structural similarity searches, we identified 25 groups with exactly 1 protein from each human herpesvirus and 89 homology groups. Many of these clusters matched these previous attempts to categorize herpesvirus proteins in groups based on experimental data and sequence analysis, confirming our approach. An example are the homologs of the alphaherpesvirus UL25 proteins, which are essential capsid-associated proteins and part of the portal cap^19^. We also found clusters of proteins with known and verified enzymatic functions, such as the herpesvirus serine/threonine kinases, named after the prototypic HSV-1 US3 and UL13 proteins (Figure 2B). As only the protein models that passed the quality scores were used, not all groups contain exactly one protein from each species. This was seen for the proliferating cell nuclear antigen (PCNA)-like protein, large tegument protein, and tegument protein 7 clusters. For example, HCMV UL44 is missing from the PCNA group because it was deemed a low-confidence model.

Surprisingly, the DNA polymerase catalytic subunit and DNA helicase/primase complex-associated protein were found to be structurally similar (Figure 2C). While these proteins have different biological functions, this clustering can be explained by partial sequence similarity in the C-terminus^23,24^. Furthermore, they are both similar to nucleases, with HSV-1 UL8 being homologous to the exonuclease domain of B-family DNA polymerases^24^ and HSV-1 UL30 having RNase H activity^25^. Since all viruses have both of these genes, this suggests that while these genes likely share a common ancestor, they have diverged in sequence and function.

The small capsomere-interacting protein (SCP) is conserved in all herpesviruses. However, they clustered as distinct groups based on the subfamily (Figure 2D). Genetic drift may have led to structurally distinct features, as the last common ancestor likely predates the divergence of the subfamilies. This was seen with both the sequence and structure based clustering.

A particularly interesting cluster of so far-unannotated proteins consisted of the likely homologs of HCMV UL112-113. HCMV UL112-113 is essential for viral replication by mediating replication compartment formation through phase separation at viral genomes, thereby recruiting the viral polymerase^16^. The poorly characterized roseolovirus U79 proteins and the alphaherpesvirus UL4/ORF56 proteins cluster with UL112-113. They shared an undescribed and conserved predicted beta-barrel (Figure 3A). These herpesviruses also form replication compartments and have a similar subcellular localization^16,26^, and their potential UL112-113 homologs would be prime candidates for further investigation.

**Figure 3.**
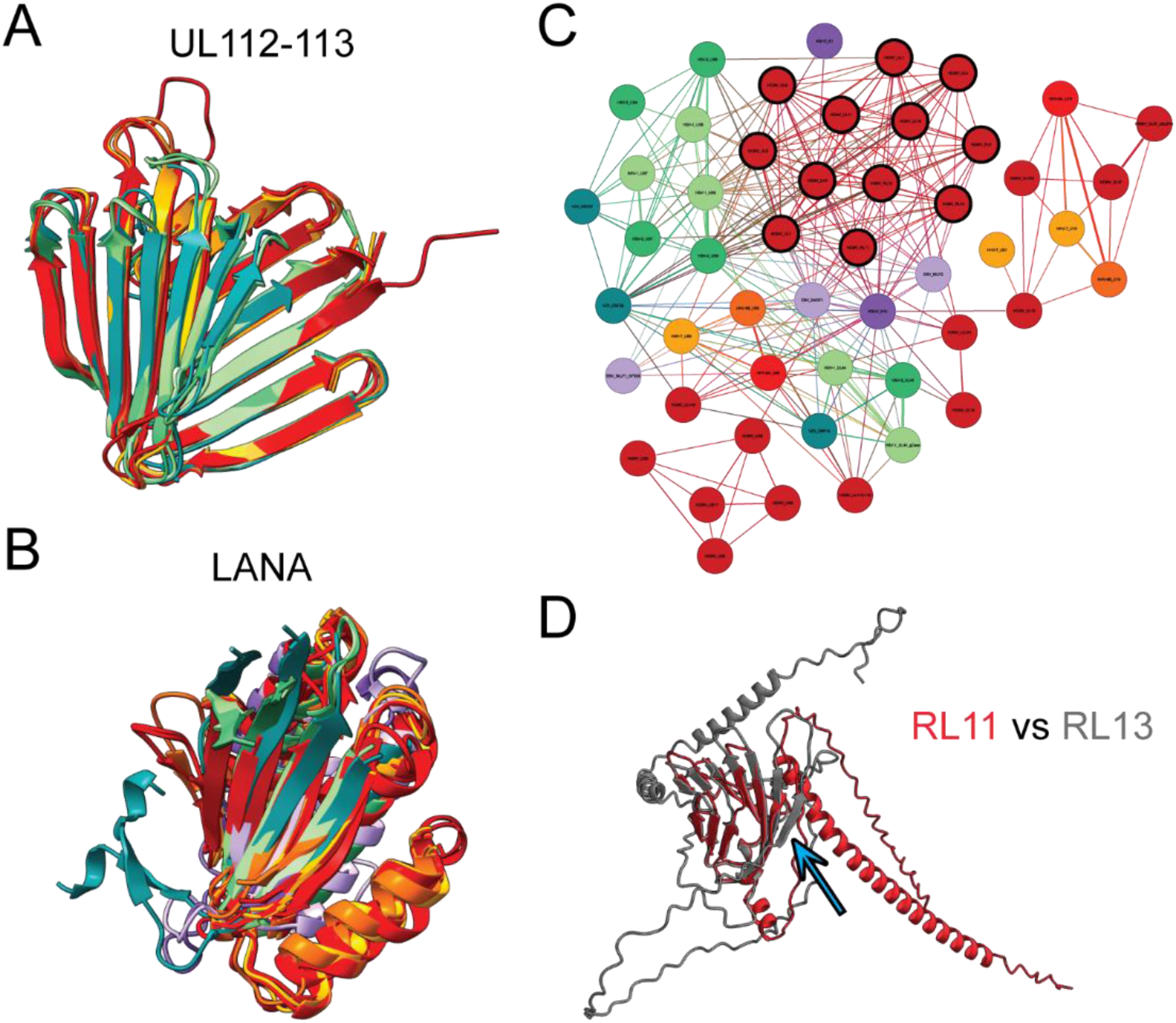
Structural clustering identifies differently grouped protein groups. A. Structural alignment of the HCMV UL112-113 cluster. Only the conserved beta-barrel-like domain is shown for clarity of HCMV UL112-113, HHV-6A U79_U80, HHV-6B U79, HHV-7 U79, HSV-1 UL4, HSV-2 UL4, and VZV ORF56. B. Structural alignment of the LANA DNA binding domain-containing cluster with only the matching domain shown for clarity of KSHV ORF73, EBV EBNA1, HHV-6A U84, HCMV UL117, HCMV UL122, HCMV UL123, HHV-6A U86, HHV-6A U90_U87U86, HHV-6B U84, HHV-6B U90_U86, HHV-7 U84, HHV-7 U86, HSV-1 UL3, HSV-2 UL3, and VZV ORF58. C. Ig-like domain-containing cluster harboring the HCMV RL11 protein family. The canonical members are marked with a black outline. D. Structural alignment of RL11 with RL13. The blue arrow points to the additional beta-strand. The corresponding UCSF ChimeraX sessions can be found at https://zenodo.org/records/13284140.

Another cluster contained the gammaherpesvirus KSHV ORF73, also known as LANA, and EBV EBNA1 proteins that harbor a known DNA binding domain^27,28^ (Figure 3B). Our structure predictions indicate that this DNA binding fold is conserved amongst all human herpesviruses, and we further analyzed these proteins by generating AlphaFold 3 predictions of dimers^29^ with DNA (Supplementary Fig. 4). The alphaherpesvirus UL3/ORF58 proteins showed similar surface charge distributions and DNA binding sites as KSHV and EBV. In contrast, the betaherpesvirus UL122 and U86 proteins, which have known IE2 transcriptional regulation activity^30,31^, bound to DNA with a different orientation in the predictions. Surprisingly, the betaherpesvirus proteins UL117 and U84 were not predicted to bind DNA, which is consistent with the less prominent positive surface charge. Perhaps these proteins have lost the DNA binding activity and acquired a different function.

The HCMV phosphoprotein UL25 formed a group with its roseolovirus U14 homologs and clustered with the HCMV UL35 proteins, which likely arose via gene duplication, constituting the pp85 superfamily^32^. Strikingly, we also found HCMV UL32 and its roseolovirus U11 homologs to be structurally similar. UL32, also known as pp150, is a protein conserved in the betaherpesviruses. It binds directly to capsids as an inner tegument protein while bridging the full virion tegument via its disordered C-terminus^33^. It will be essential to dissect if the structurally similar UL25 and UL35 can also bind capsids and potentially regulate pp150 abundance at pentons, potentially confusing the reconstruction of virus capsid cryo-EM maps. Finally, the poorly studied KSHV ORF48 and likely the EBV BBRF2 protein, which was excluded from the clustering due to low prediction scores, also clustered with the pp150 homologs. ORF48 is part of virions^34^, and it should be assessed if it can bind to the viral capsid as pp150 does.

Since some protein predictions have unstructured regions, it is possible that these flexible regions lead to false negative results by Foldseek. To evaluate this possibility, all predicted structures were cut into smaller pieces based on where they are likely structured, *i.e.* pLDDT >0.7. Foldseek clustering was then repeated and compared to clustering based on full-length predictions (Supplementary Data 2 Sheet 9). While this method resulted in an interesting novel cluster containing EBV BALF1, EBV BHRF1, and KSHV ORF16, which are all known apoptosis regulators, most novel clusters identified using this method were likely false-positives such as HSV-1 UL49 and UL49A. UL49 is a tegument protein involved in immune evasion, and UL49A is a glycoprotein, respectively. Another example is a cluster containing HCMV UL96 and HSV-1 UL14, which show little unique or similar tertiary structure. For this reason, we focused on the analysis of full-length proteins.

We also compared our results to sequence-based clustering by MMseqs2 and HHblits (Supplementary Data 2 Sheet 2 and 3, respectively). Importantly, several structure-based clusters contained additional proteins when compared to sequence-based clustering methods, such as the UL112-113 cluster and the KSHV ORF73 (LANA) cluster, where all the identified proteins of this cluster have functions related to DNA binding. However, we also identified cases where structure predictions failed the quality scores and HHblits-derived clusters were more complete. For example, all pAP proteins, which are splice variants of the scaffold protein, failed the structure prediction quality scores and are absent from the structural cluster. Since sequence-based methods can complement these instances where the structure prediction is of low quality, we provide an option to view both the structure and sequence-based clusters at https://www.herpesfolds.org/herpesclusters. Supplementary Data 2 Sheet 10 lists the respective HHblits and Foldseek scores for each protein pair as well as all interactions above their respective thresholds, which were found only by HHblits, only Foldseek, or both.

### Immunoglobulin-like domains are common structural elements

One of the largest clusters we identified using structural clustering via Foldseek consists of viral proteins that contain immunoglobulin (Ig)-like domains. This cluster merged several previously defined protein families. One of the included groups is the HCMV RL11 family^35^. They share an RL11 domain (RL11D), formed by a conserved tryptophan and two cysteines, and are structurally similar to immunoglobulin domains. The RL11 proteins are non-essential in cell culture^36^ but have high sequence variability between isolates with some playing a role in modulating immune responses, such as UL11^37^. While there are 14 accepted members of this family, they did not form an isolated structural group in our clustering (Figure 3C). Unexpectedly, EBV BILF2 and KSHV K14 but also the well-studied HSV-1 glycoproteins UL44 (gC), US6 (gD), US7 (gI), and US8 (gE) were structurally similar to multiple members of this family. EBV BILF2 is encoded in a genome region unique to *lymphocryptoviruses*, and KSHV K14 is also unique to KSHV. Furthermore, HSV-1 and HSV-2 US8 are located in a different part of the genome than HCMV. In addition to the low sequence identity, these data suggest that these Ig-like domain-containing proteins were acquired independently rather than from a common ancestor. Furthermore, gene duplication continued after speciation. While most RL11 members have the same 6-strand antiparallel beta-fold, RL13 has an additional beta-strand (Figure 3D), potentially representing a distinct evolutionary branch and function.

### Identification of domain-level duplications and deletions

We performed the initial structural similarity search with full-length proteins. While this will identify commonalities between proteins, it will, by definition, not identify differences. To address how the domain architecture of a protein can differ between species, we performed a structural similarity search with overlapping structural snippets using a sliding window approach (see materials and methods) (Supplementary Data 2 Sheet 6). We identified 2806 structurally similar connections between proteins, which includes 183 new connections for proteins relative to the full-length analysis (Supplementary Data 3 Sheet 1 and 2). This analysis identified 24 internal duplications, where domains were likely duplicated in the same protein; 35 repetitive acquisitions, where domains of a different protein matched the query protein multiple times at different positions; and 13 domain additions, where multiple domains of a protein matched different proteins (Figure 4A and Supplementary Data 3 Sheet 3).

**Figure 4.**
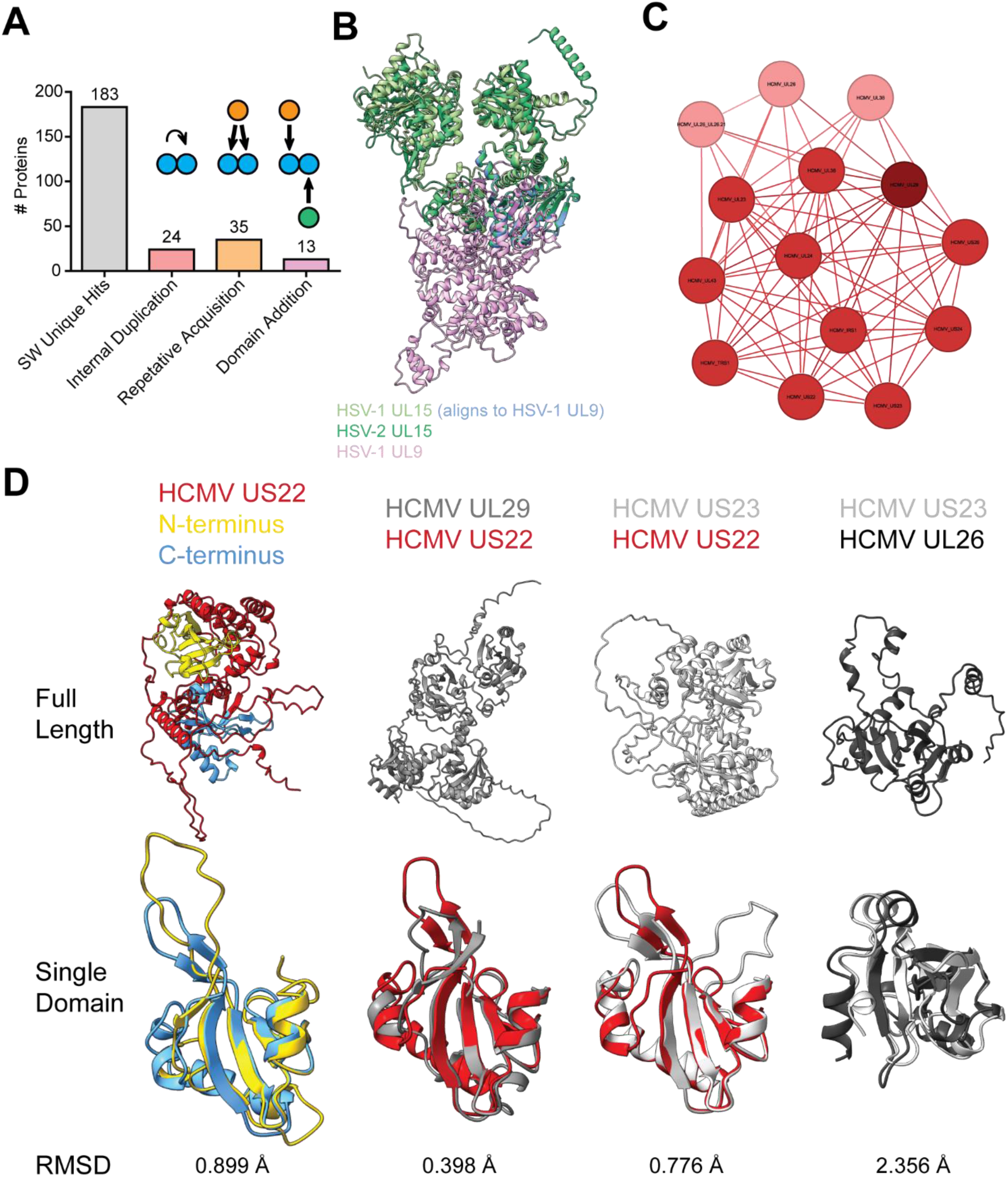
Domain-level similarity search identifies conserved domain duplications. A. Domain-level similarities identified using a sliding window approach. The "SW Unique Hits" are the number of proteins that had a structurally similar query-target pair in the sliding window analysis that was absent from the full-length analysis. "Internal Duplications" are proteins where parts of the protein are similar to another part of itself. "Repetitive Acquisition" are proteins where a piece of a different protein matches the query protein multiple times at different positions. "Domain Addition" refers to proteins with multiple domains matching different proteins. Illustrative cartoons are shown above the corresponding bar. B. HSV-1 UL15 is aligned to the known homolog HSV-2 UL15 and the structurally similar target HSV-1 UL9, which was only identified in the domain-level search. The part of HSV-1 UL15 that aligns with HSV-1 UL9 is colored blue. C. The cluster of HCMV proteins that contain HCMV US22. The proteins that contain 2 domains, like the canonical US22, are shown in the HCMV color. The proteins with 1 domain are shown in light red, and the protein that has 4 domains is shown in dark red. D. The tertiary structure of the core domain is similar between the proteins in this group. The top row is the full-length protein, the bottom row is a structural alignment of the core domain, and the RMSD against the 91 Cα of the core domain alignment is written below. HCMV US22 has 2 domains, shown in yellow and blue, which are aligned. HCMV UL29 has 4 domains and was aligned to US22. HCMV US22 was aligned to HCMV US23, which has 2 domains, and HCMV US23 was aligned to HCMV UL26, which has 1 domain. In the case of multiple domains, the best-fitting domain combination is shown. The corresponding UCSF ChimeraX sessions can be found at https://zenodo.org/records/13284140.

We found that HSV-1 UL15 was structurally similar to the previously identified HSV-2 UL15 but also HSV-1 UL9 (Figure 4B). UL15 is part of the tripartite terminase, essential for packaging viral genomes into newly formed capsids, and UL9 is the replication origin-binding protein. The fragment that aligns is oriented in the opposite direction relative to the core in UL15 versus UL9, which suggests that this region is not involved in an external interaction and has a conserved structural integrity function.

An important example of domain duplications is in the large betaherpesvirus US22 family cluster. No experimental structure of this large family of proteins is available, and sequence-based methods indicate that the family consists of at least 12 members in HCMV^38,39^. Using full-length structural similarity searches, we identified HCMV proteins that share the same conserved fold (Figure 4C). Importantly, this group consists of four different domain architectures depending on whether the protein contains 0, 1, 2, or 4 copies of the US22 domain (Figure 4D). The majority of proteins are similar to US22 with two domains, while the following members have only one domain: HCMV UL26, HCMV UL26_UL26.21, HCMV UL27, HCMV UL38, HHV-6A U19, HHV-6B U19, and HHV 7 U19. We also found a subgroup that lack a US22 homology domain consisting of HHV-6A U4, HHV-6B U4, and HHV-7 U4. They are connected to the main group through the HHV-6A U7 and HHV-7 U7 proteins, which appear to have an additional domain relative to US22, which the U4 proteins consist of exclusively. The proteins with four copies of the US22 domain are HCMV UL29, HHV-6A U7, and HHV-7 U7. Oddly, HHV-6B U7, unlike the other roseoloviruses, only has two domains, suggesting a divergence in this species. Furthermore, many group members were identified to contain an internal duplication and/or repetitive acquisition. The low RMSDs between the two domains of US22 and between the domains from different proteins are consistent with a domain duplication (Figure 4D). These data suggest that the US22 family is bigger than previously thought and that domain additions and deletions are particularly common in this family.

### Viral exaptation of cellular proteins

Many of the identified clusters contained understudied proteins. We expanded the structural similarity search to cellular proteins using Foldseek to determine their potential biological activity from the PDB database (Supplementary Data 4 Sheet 1), and the predicted AlphaFold structures of the Swiss-Prot database (Supplementary Data 4 Sheet 3). The PDB contains experimentally verified structures, but it is limited in diversity. In contrast, the AlphaFold predictions of the Swiss-Prot database enable comparisons to a much more extensive predicted database that is less biased by experimental limitations. We combined the significant hits from the two databases and identified structurally similar cellular proteins. To find putative functions for the viral protein groups, the functional description of the homologous proteins was mined for keywords (Supplementary Data 4 Sheets 2 and 4). Unfortunately, the depositors’ descriptions of structures in the PDB database are not systematic. To handle these variations, we calculated the frequencies of all words in the descriptions, and the words with the highest frequencies, which likely represent the protein function, were manually chosen as the annotation. Each viral homology group identified above was then given a HerpesFolds annotation. Since the PDB, but not the AlphaFold database, contains structures of viral proteins, it is important to note that some hits in the PDB database are because they were aligning with themselves.

Some predicted viral protein structures did not map to any other protein fold in the searched databases, indicating that these are unique folds (Supplementary Data 4 Sheet 5). We found that even though the viral alkaline nucleases are not similar to cellular nucleases, they are consistent within the herpesviruses (Figure 5A). Experimental structures have been solved for EBV^40^ and KSHV^41,42^. While the central core was found to show some similarity to λ-exonuclease, the viral proteins have additional terminal extensions. Another striking example was the cytoplasmic envelopment protein 2, which is conserved in all herpesviruses and for which no experimental structure exists (Figure 5A). Still, this protein group was very well predicted as indicated by its pLDDT and PAE scores. Moreover, the most distant phylogenetic pair, VZV ORF44 and HHV-6B U65, had an RMSD of 4.3 Å against the 112 Cα atoms despite a sequence identity of only 14%. These results indicate that AlphaFold can predict herpesvirus proteins with high confidence *de novo* even if no structurally similar dataset exists. Since the CEP2 proteins are crucially involved in virion envelopment^43^, it will be important to analyze their role in modulating the cellular secretory system through this unique fold.

**Figure 5.**
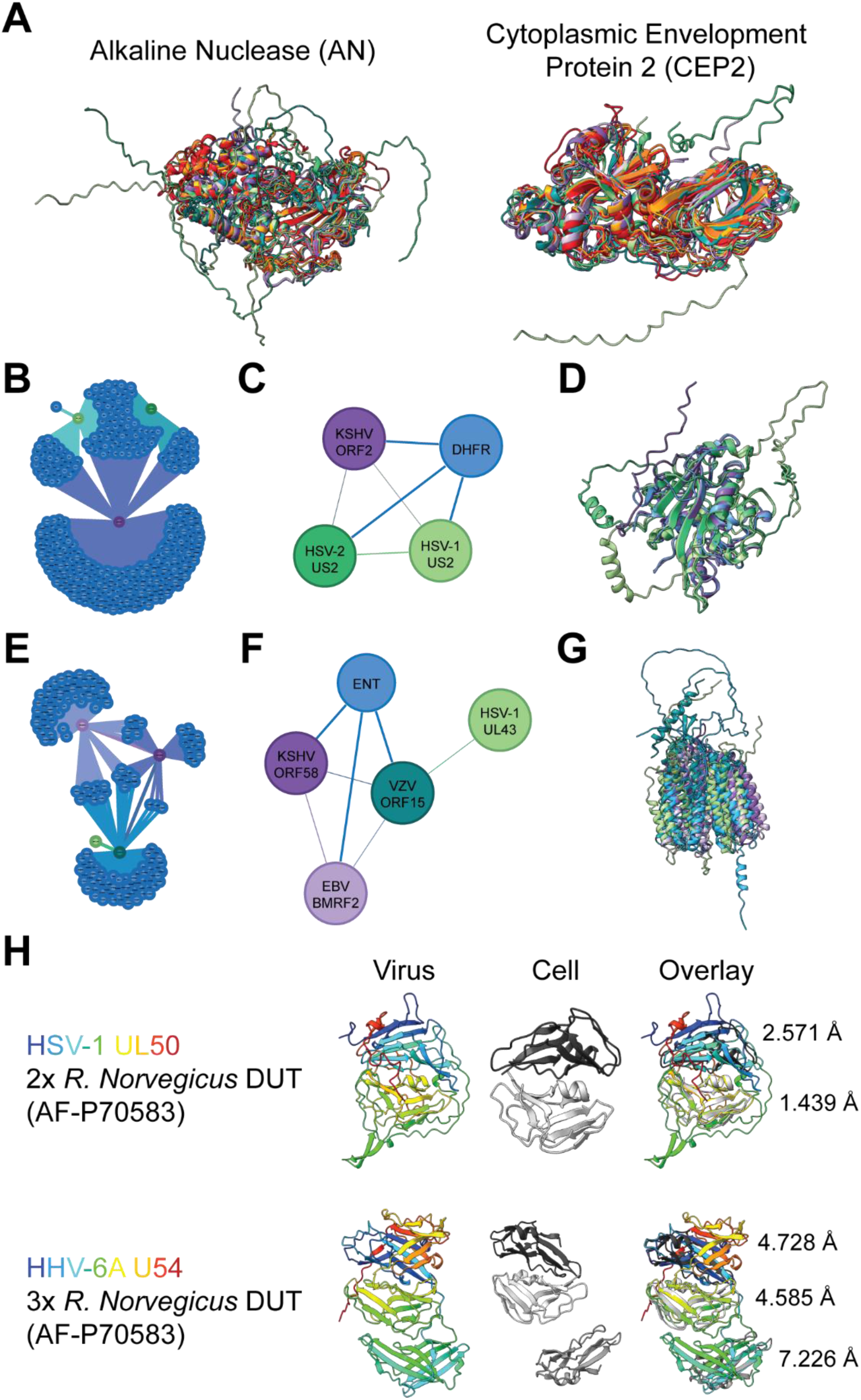
Annotation of viral protein functions from structural similarity search. A. Structural alignment of the alkaline nuclease (AN) and cytoplasmic envelopment protein 2 (CEP2) from all human herpesviruses. CEP2 was found in all human herpesviruses but had no significant cellular hit. B. Cluster of KSHV ORF2, HSV-1 US2, and HSV-2 US2 containing the cellular hits. Each cellular node (blue) represents a specific protein. C. Summary of B for clarity. D. Structural alignment of the proteins in C. P04753 dihydrofolate reductase (DHFR) was the most significant cellular hit. E. Cluster of EBV BMRF2, KSHV ORF58, HSV-1 UL43, and VZV ORF15 containing the cellular hits. Each cellular node (blue) represents a specific protein. F. Summary of E for clarity. G. Structural alignment of the proteins in F. The most significant cellular hit was Q8R139 equilibrative nucleoside transporter 4 (ENT4). H. Structural architecture of the classical DUT HSV-1 UL50 and the DURP HHV-6A U54. The rainbow-colored viral protein illustrates how the polypeptide folds back on itself. The most significant cellular hit, AF-P70583 DUT from *Rattus norvegicus*, is shown multiple times so that it can be aligned to each domain in the viral protein. The RMSD against the 205 Cα for each alignment is shown on the right of the overlay. The disordered termini were hidden for clarity. The corresponding UCSF ChimeraX sessions can be found at https://zenodo.org/records/13284140.

Using this workflow, we could annotate several poorly studied viral proteins and identify potential functions. One example is HSV-1 US2, which is structurally similar to dihydrofolate reductases (DHFRs) (Figure 5B-D). The most significant cellular target was the DHFR from *Mesocricetus auratus* (commonly known as the golden or Syrian hamster) (PDB 3EIG^44^). Furthermore, HSV-1 US2 was structurally similar to KSHV ORF2, a known DHFR homolog with enzymatic activity^45,46^. While KSHV ORF2 contains the catalytic residues identified in *E. coli*^47,48^, HSV-1 and HSV-2 US2 do not contain them (Supplementary Fig. 5). Since no DHFR activity has been reported for the US2 proteins, this fold was likely exaptated. Functional assays will be needed to verify if the US2 proteins still have DHFR activity.

We also identified a cluster containing EBV BMRF2, KSHV ORF58, VZV ORF15, and HSV-1 UL43 (with HSV-2 UL43 being absent due to low-quality scores). Importantly, EBV BMRF2, KSHV ORF58, and VZV ORF15 predictions were significantly similar to the human equilibrative nucleoside transporter (ENT) 4 (Figure 5E-G). This cellular protein is involved in transporting metabolites and regulating adenosine concentrations^49^. However, no experimental data currently supports a role for any of these viral proteins in transmembrane transport. Instead, EBV BMRF2 can change cellular morphology by modulating the actin cytoskeleton^50^ and is essential for efficient cell-cell spread^51^. In a closely related alphaherpesvirus, UL43 colocalizes with proteins involved in membrane fusion and potentially interacts with the gM/gN complex^52^, which is engaged in syncytia formation^53^. These commonalities suggest a function for this viral protein cluster in cell-cell spread, which does not align with the transmembrane transporter activity of the structurally similar cellular proteins. Moreover, comparative cross-sections of the viral proteins and ENT4 illustrate that they do not have a channel necessary for metabolite transport^54^ (Supplementary Fig. 6A). Several transporters depend on accessory proteins for correct trafficking to the plasma membrane, such as the lysosomal transporter MFSD1, which depends on GLMP^55^. This is also true for EBV BMRF2, which needs BDLF2^50^ for correct trafficking to the plasma membrane, and KSHV ORF58, which needs ORF27^34^. Interestingly, AlphaFold predictions of the viral heterodimers suggest that even if channels were present in BMRF2 and ORF58, they would be blocked by the partner protein (Supplementary Fig. 6B), arguing against a conserved transporter activity as recently proposed^56^. Instead, the structure predictions likely indicate that the similarity to ENT4 may stem from "radical" exaptation, where the transporter activity was lost and replaced by a role in viral cell-to-cell spread.

All human herpesviruses encode for deoxyuridine triphosphatases (dUTPases)^57^ called HSV-1 UL50, HSV-2 UL50, VZV ORF8, HCMV UL72, U45 of roseoloviruses, KSHV ORF54 and EBV BLLF3. Cellular dUTPases are homotrimers that form three active centers at their interfaces. In contrast, the herpesvirus dUTPases are monomers consisting of a fusion of two dUTPase domains, as shown for EBV BLLF3^58^. While the respective alpha-and gammaherpesvirus proteins still have enzymatic activity, the betaherpesvirus homologs have lost them and are involved in immune modulation^59^. Human beta-and gammaherpesviruses code for 17 more dUTPase-related proteins (DURPs), likely due to gene duplications^57^. Using domain-level search, we found that all DURPs were predicted to harbor a three-domain architecture in contrast to the two domains of the viral dUTPases. However, the separate domains diverged quite significantly from the canonical dUTPase fold as indicated by their RMSD (Figure 5H), which is in line with a recent report on the crystal structure of the HCMV DURP UL82 (pp71), which describes that it has lost its dUTPase function^60^.

Herpesviruses have been suggested to induce autoimmunity through molecular mimicry^61,62^. Several studies have attempted to identify viral proteins that might induce autoimmunity due to a similarity to a cellular protein^63,64^. Protein sequence-based analyses could not identify viral proteins that would induce autoimmunity to known cellular proteins^63^, such as myelin for multiple sclerosis^64^. Our structural similarity search identified no direct structural match of myelin to a viral protein. Interestingly, multiple members of the immunoglobulin domain-containing group, such as HSV-1 US8, mapped to myelin protein zero-like protein 1, and others, such as EBV BARF1, mapped to myelin-associated glycoprotein. These myelin-related proteins could be involved and warrant further investigation.

## Discussion

Understanding protein structures is critical to comprehending pathogen-host interactions. Unfortunately, experimental structural data is lacking for most human herpesvirus proteins. Machine learning-based structure prediction tools have made huge strides in accuracy and provide quality scores for their predictions^1,2,11^. Accordingly, recent community evaluations have found "that AF2 [AlphaFold2]-predicted structures, on average, tend to give results that are as good as those derived from experimental structures"^3^ while also cautioning that "AlphaFold predictions are valuable hypotheses and accelerate but do not replace experimental structure determination"^12^.

The human herpesviruses encode for at least 844 proteins, and for most, no experimental structure has been solved. Here, we applied proteome-wide predictions to provide a structural systems view of all annotated human herpesvirus proteins and developed stringent scoring workflows to validate our predictions. We generated clusters of proteins with similar folds using all-versus-all full-length structural similarity searches. Importantly, we identified previously unknown protein clusters sharing conserved folds, such as the UL112-113 family that carries a beta-barrel-like fold and likely plays an essential role in genome replication, while confirming previously known clusters, such as viral G-protein coupled receptors and kinases. In comparison, previous work has used sequence-based approaches to cluster herpesvirus proteins into related families. Importantly, neither a sequence-based alignment algorithm (MMseqs2) nor an HMM-HMM-based algorithm (HHblits) could uncover all of the novel and biologically important relationships that we found using a structural approach, such as the finding that the alphaherpesvirus UL3 proteins are new members of the KSHV ORF73 (LANA) cluster, which are DNA-binding proteins and have too little sequence conservation to be identified by MMseqs2 or HHblits.

Still, structural clustering might not always indicate direct evolutionary relatedness. The large Ig-like domain cluster appears to be a group of proteins with a common fold, likely the result of independent acquisitions rather than originating from a single common ancestor or gene duplication. Using domain-level searches, we identified differences between structurally related proteins such as viral dUTPases and the US22 family. Both consist of subgroups with different numbers of duplicated domains that correlate with their described activity. Finally, we searched for cellular proteins with matching folds. We identified cases of potential exaptation, such as the membrane transporter ENT4 by human alpha-and gammaherpesviruses and unique folds that do not match any cellular protein, such as the CEP2 proteins.

This body of data provides many valuable hypotheses for experimental validation. To accelerate the use of this data and research on human herpesvirus, we have generated an easy-to-use, searchable web interface for the community, accessible at https://www.herpesfolds.org/herpesfolds. The database groups proteins by structural relationship and allows interactive evaluation of all structures. PDB files of each prediction can be downloaded or directly submitted to the Foldseek structural similarity search server to find similar cellular proteins in up-to-date databases.

This work is a first step based on high-confidence monomeric predictions of viral proteins. It neglects that some proteins may fold differently in complex with others, adopt multiple conformations, or need to form oligomers to function, such as gB^65^. Future work will need to take into account protein complex composition and stoichiometry. However, accomplishing this task is challenging as this information is unavailable for most viral proteins and generating all likely and possible combinations is computationally highly demanding. The next logical step will be predicting viral protein-protein interaction networks. Since post-translational modifications and protein-ligand complexes^66,67^ or nucleic acids^68^ will likely play a role in many such complexes, the recently released RoseTTAFold-All-Atom^69^ and AlphaFold 3^70^ algorithms will be of great use.

Herpesviruses have large bicistronic genomes, and for example, ribosomal profiling has identified 100s of additional transcripts and ORFs for HSV-1^71^, HCMV^72^, EBV^73^ and KSHV^74,75^. Structure predictions of these could provide insights into their functions and roles during infection and pathogenesis and should be the next step.

The methodology presented here applies to any virus or virus family. It can easily be extended to analyze all reading frames of a given organism, not only the annotated ORFs. As the databases of predicted and experimental viral protein structures grow, more connections between different viral pathogens and their hosts will be made. The speed at which these *in silico* models can be generated will drive rapid hypothesis design and subsequent experimental confirmation.

## Methods

### Viruses

The strains and associated reference numbers of the nine human herpesvirus proteomes used for prediction are noted in Table 1. In addition to the full-length proteins, splice variants were also included. A complete list of the predicted proteins can be found in Supplementary Data 1 Sheet 4.

### Software

Sequence alignments were performed with MMseqs2 release 15-6f452^76^ (https://github.com/soedinglab/MMseqs2) and HHblits v3.3.0^77^ (https://github.com/soedinglab/hh-suite). HHblits HMM generation used the UniRef30 database from 2023_02. The significance threshold used for MMseqs2 was E-value <0.1^76^ and for HHblits was E-value <0.001 (https://toolkit.tuebingen.mpg.de/tools/hhblits). Protein structure predictions were made with LocalColabFold v1.5.1^11^ (https://github.com/YoshitakaMo/localcolabfold) and AlphaFold^1^ using AlphaFold version 2.3.0 (https://github.com/google-deepmind/alphafold). Fasta files were downloaded from https://www.uniprot.org, and structure predictions were performed in accordance with the respective documentation detailed in the respective GitHub releases. For each protein, 5 models were generated using 3 recycles and a stop-at-score of 100. Proteins that failed the quality scores were rerun with LocalColabFold with 20 recycles. AlphaFold 3^70^ predictions were performed through the AlphaFold Server at https://alphafoldserver.com. Protein topology was identified with DeepTMHMM^78^ using their provided Demo Colab (https://dtu.biolib.com/DeepTMHMM). The custom python script “DeepTMHMM_export_domains.py” consolidated the DeepTMHMM outputs into a single file and “DeepTMHMM_make_truncated_fasta.py” generated fasta files of the proteins without the identified signal peptide. Structural similarity searches were performed with local install versions of DALI^22^ (DaliLite.v5 from http://ekhidna2.biocenter.helsinki.fi/dali/) and Foldseek^5^ version 2ad017897d3dab66dd33ea675e92215bdfb4a64d (https://github.com/steineggerlab/foldseek). Networks were visualized with Gephi v0.10.1 (http://gephi.org/). UCSF ChimeraX version 1.8 (https://www.cgl.ucsf.edu/chimerax) was used for visualization of structures, and PyMOL 2.5.8 (https://www.pymol.org) was used for root mean square deviation (RMSD) calculations. Data analysis was carried out with custom-written Python scripts, which were partly generated with the assistance of large language models and thoroughly tested manually. The complete set of scripts can be accessed through the project’s GitHub repository: https://github.com/QuantitativeVirology/Herpesfolds.

### Evaluation of model quality

All proteins were initially predicted using LocalColabFold^11^ with 3 recycles as it is more time-efficient. To assess the quality of the resulting models, 3 thresholds were used. Failing any single threshold led to the model being flagged as "fail". We interrogated whether the model optimization had converged by testing the consistency of the predicted local distance difference test (pLDDT) scores associated with every model (Supplementary Fig. 1A and B). To do so, we first calculated the mean and standard deviation of the pLDDT values per model and then the standard deviation of these 5 values, resulting in “StDev of mean” and “StDev of StDev”. To set thresholds, we fit a Gaussian curve to their respective histograms and set the cutoff at the mean of the Gaussian plus 2 times its standard deviation (Supplementary Fig. 1C and D). Therefore, a “StDev of mean” >3.2 or a “StDev of StDev” >1.9 was deemed a low-quality model. We also evaluated the predicted template modeling score (pTM) to validate model confidence further. To set a global threshold at which a prediction likely constitutes a folded protein, we chose 106 herpesvirus proteins representing the protein length distribution of the herpesvirus proteomes and scrambled their amino acid sequences. These scrambled sequences were used for model prediction and treated as a negative dataset, assuming that a random residue order should not fold. Receiver Operating Characteristic (ROC) analysis yielded an area under the curve of 91.3%, and the Youden index was used to set a threshold (Supplementary Fig. 1E). Following this analysis, a pTM <0.3150 was deemed a low-quality model. We reanalyzed all models that failed the initial threshold in AlphaFold^1^ and LocalColabFold^11^ with 20 recycles and rescored them. For example, Supplementary Fig. 1F shows an improved model run in AlphaFold with higher pLDDT scores compared to Supplementary Fig. 1B. The corresponding scripts can be found in our HerpesFolds GitHub repository in the file “score_colabfold.py” at https://github.com/QuantitativeVirology/Herpesfolds.

### Defining disordered and structured content of models

The pLDDT value at a given residue indicates whether a region is predicted to be disordered (pLDDT <0.50) or structured (pLDDT >0.70). For the models that passed all quality scores, we generated histograms of the percentage of the protein below or above these values and fit a Gaussian to both curves. Since the majority of proteins are expected to be structured, a threshold, >44%, for proteins containing a disordered region was set at the mean of the distribution + 2x StDev for the percentage of protein sequence below 0.50, and a threshold, >33%, for proteins with a structured region was set at the mean -2x StDev for the percentage of protein sequence above a pLDDT of 0.70. The script for extracting pLDDT values from the .json files associated with every prediction can be found in our HerpesFolds GitHub repository in the file “score_colabfold.py” at https://github.com/QuantitativeVirology/Herpesfolds.

### Extracting pLDDT domains from predicted structures

The structured domains from a predicted structure were extracted based on the pLDDT scores. The regions with a pLDDT >0.7 were identified. If adjacent regions were connected by a disordered region, *i.e.* pLDDT <0.7, that was shorter than 100 residues, the adjacent regions along with the disordered region were counted as a single domain. The corresponding script can be found as “extract_pLDDT_domains.py” in our HerpesFolds GitHub repository at https://github.com/QuantitativeVirology/Herpesfolds.

### Structural similarity search with DALI and Foldseek

Foldseek aligns protein structures with decreased computation time compared to alternative algorithms^5^. It provides different alignment algorithms, 3Di+AA Gotoh-Smith-Waterman and Tmalign, and measures of significance, such as the Expect-value (E-value) and probability of homology (prob). We tested both alignment algorithms with different thresholds and compared the protein clusters, Supplementary Data 2 Sheet 7. The Tmalign results did not agree with the sequence based clustering and known literature. The E-value and prob thresholds with 3Di+AA Gotoh-Smith-Waterman gave similar results, and thus we chose to continue with an E-value threshold of 0.001 to be consistent with the literature^10^, Supplementary Data 2 Sheet 8.

To cluster viral proteins into groups of structural similarity, we employed DALI as an established algorithm and Foldseek as a recent algorithm designed to work with vast predicted datasets with decreased computation times. With DALI, we used the recommended threshold of Z-score >8 for probable homologs^79^, and with Foldseek, we used the previously published threshold of <0.001^10^. Since the results from both algorithms were similar, we only used Foldseek for virus-host and domain-level searches as DALI is orders of magnitude slower, which would have made these analyses computationally too expensive otherwise. Similarity searches against cellular proteins were done using the PDB database (Supplementary Data 4 Sheet 1), and the predicted AlphaFold structures of the Swiss-Prot database (Supplementary Data 4 Sheet 3).

### Clustering of structure predictions by similarity

The Foldseek and DALI outputs are a list of query and target proteins with the associated significance value. We used a threshold of E-value <0.001 for Foldseek and a Z-score >8 for DALI to determine the significant results. Both reciprocal comparisons for each protein pair needed to be significant for inclusion in the clustering, *i.e.* A matches B and B matches A. If protein A was structurally similar to B and B similar to C, then A, B, and C were clustered. Once all clusters were identified, we converted the protein names to their row position in Supplementary Data 2 Sheet 1. This system allowed us to quickly determine whether a structural cluster contained proteins expected to group, *i.e.* all row numbers are the same, or not, *i.e.* the row numbers are different, based on UniProtKB annotations^20^ and previously known homologies^21^.

The associated script can be found in our HerpesFolds GitHub repository in the file “analyze_foldseek_for_structural_similarity_cluster.py” at https://github.com/QuantitativeVirology/Herpesfolds.

### Analysis of sliding window structural similarity

To search for alterations at the domain level between protein predictions, we generated “structural snippets” from all predictions using a sliding window approach. We then ran these structural snippets in an all-versus-all Foldseek search to identify conserved folds. We empirically tested which window size would result in the highest number of significant hits using Foldseek. Trying a window size of 25, 50, 100, 150, and 200 amino acids, we found the most significant hits with a window size of 150. The sliding window step size was tested at 5, 10, 15, 20, and 25 amino acids. We used a step size of 20 amino acids for the final analysis as it resulted in the maximal number of protein species detected. This approach resulted in over 12000 structural snippets of 150 residues in length, each overlapping by 130 residues with the preceding fragment and covering the whole folded human herpesvirus proteome. All structural fragments were compared pairwise with Foldseek, accounting for about 1.6 million tested combinations. We next assembled “structural contigs” of overlapping frames that passed the Foldseek threshold of an E-value <0.001. If a structural snippet matched another structural snippet and they were not overlapping in the same original protein, then the pair was deemed a “hit”. If several structural snippets overlapped in sequential order, they were used to build larger domains. This analysis resulted in a list of structured areas, which we used to determine if a query protein has a different domain architecture than a structurally similar protein in this data set. If a snippet of a query protein matched a different part of the query protein, we deemed it to have an "internal duplication". If a target snippet matched the query protein at multiple locations, we considered the query protein to have a "repetitive acquisition". If the query protein was structurally similar to a target protein but the query protein had at least 1 domain that was not found in the target protein, we deemed the protein to have a “domain addition”. The associated script can be found in our HerpesFolds GitHub repository in the files “make_sliding_window_pdb.py”, “Sliding_Window_Analysis.ipynb”,“analyze_foldseek_sliding_window.py”, and “analyze_sliding_window_domains.py” at https://github.com/QuantitativeVirology/Herpesfolds.

### Keyword analysis of Foldseek results

Most Foldseek searches against cellular databases resulted in multiple significant matches, each containing protein descriptions and matching protein identifiers. We filtered for the most enriched keywords to extract likely biological functions from these descriptions. The following terms, in addition to single characters, single and double digits, and punctuation, were filtered out as they are too generic:

crystal, a, after, allosteric, allosteric, alpha, an, and, angstrom, angstroms, antibody, apo, as, at, atomic, based, basic, beta, between, bound, by, by, c1, c-c, cell, chain, chi, class, class, coli, complex, complexed, complexes., compound, conserved, cryoem, cryo-em, crystal, c-terminal, c-x-c, de, dehydration, deletion, delta, design, different, discovery, domain, e., e.coli, edition, element, engineered, epsilon, escherichia, eta, fab, for, form, fragment, from, full, full-length, functional, gamma, herpes, herpesvirus, high, holoenzyme, ii, iii, implications, in, inhibitor, inhibitors, ion, iota, its, kappa, lambda, ligand, ligand-binding, loader, loop, low-ph, mode, molecule, monoclonal, motif, mu, mutant, natural, neutralizing, novel, novo, nu, of, omega, omicron, on, one, open, open, opposite, peptide, phi, pi, probable, protein, psi, region, resolution, reveal, rho, sigma, significance, site, specificity, structural, structure, structures, substrate, symmetry, tau, template-primer, the, the, therapeutic, theta, to, type, uncharacterized, unit, upsilon, using, variant, virus, with, xfel, xi, x-ray, zeta The used script can be found in our HerpesFolds GitHub repository at https://github.com/QuantitativeVirology/Herpesfolds in the files “Keyword_Frequency_SP_PDB.ipynb”. This script was written with the assistance of a large language model.

## Supporting information

Supplementary_Data_all

## Data Availability

The structure predictions and ChimeraX sessions generated in this study have been deposited in the zenodo database under DOI 10.5281/zenodo.13284140 at https://zenodo.org/records/13284140. Moreover, predictions and interactive clustering are available through a web interface at https://www.herpesfolds.org/herpesfolds and https://www.herpesfolds.org/herpesclusters.

## Code Availability

The scripts are available at https://github.com/QuantitativeVirology/Herpesfolds, and a permanent reference is under DOI 10.5281/zenodo.13991336 at https://zenodo.org/records/13991336.

## Acknowledgements

We would like to thank the Topf lab at CSSB for help in the initial phases of this project, Milot Midarta and Martin Steinegger for help with Foldseek and for providing links to Foldseek in HerpesFolds and Lisa Holm for advice in using DALI as a local installation. Finally, we thank Christian Löw for his help in evaluating if the BMRF2 protein cluster codes for functional transporters. This research was partly supported by the Maxwell computational resources operated at Deutsches Elektronen-Synchrotron DESY, Hamburg, Germany.

Work in the Bosse lab was funded by the Deutsche Forschungsgemeinschaft (DFG, German Research Foundation) under Germany’s Excellence Strategy EXC 2155—project no. 390874280, the DFG-funded RTG 2771 Humans and Microbes, project no. 453548970 (Bosse), the DFG-funded RTG 2887, project number 49735088, by the Wellcome Trust through a Collaborative Award (209250/Z/17/Z) and the Leibniz ScienceCampus InterACt, funded by the BWFGB Hamburg and the Leibniz Association (W75/2022) InterACt and "Hamburg-X Infektionsforschung". Moreover, the Bosse and Kaufer labs are both funded through the DFG Research Unit FOR5200 DEEP-DV (443644894) project BO 4158/5-1 and KA 3492/12-1 awarded to JBB and BBK respectively. BKK is also funded by the ERC consolidator grant (ERC-CoG ENDo-HERPES, 101087480).

## Author Contributions Statement

TKS: Conceptualization, Data curation, Formal Analysis, Investigation, Methodology, Software, Supervision, Validation, Visualization, Writing – original draft, Writing – review & editing

SO: Data curation, Software

SS: Data curation, Software

RS: Data curation, Software

BK: Data curation, Writing – original draft, Writing – review & editing, Funding acquisition, Project administration

JBB: Conceptualization, Data curation, Funding acquisition, Methodology Project administration, Supervision, Writing – original draft, Writing – review & editing

---

**Supplementary Fig. 1.**
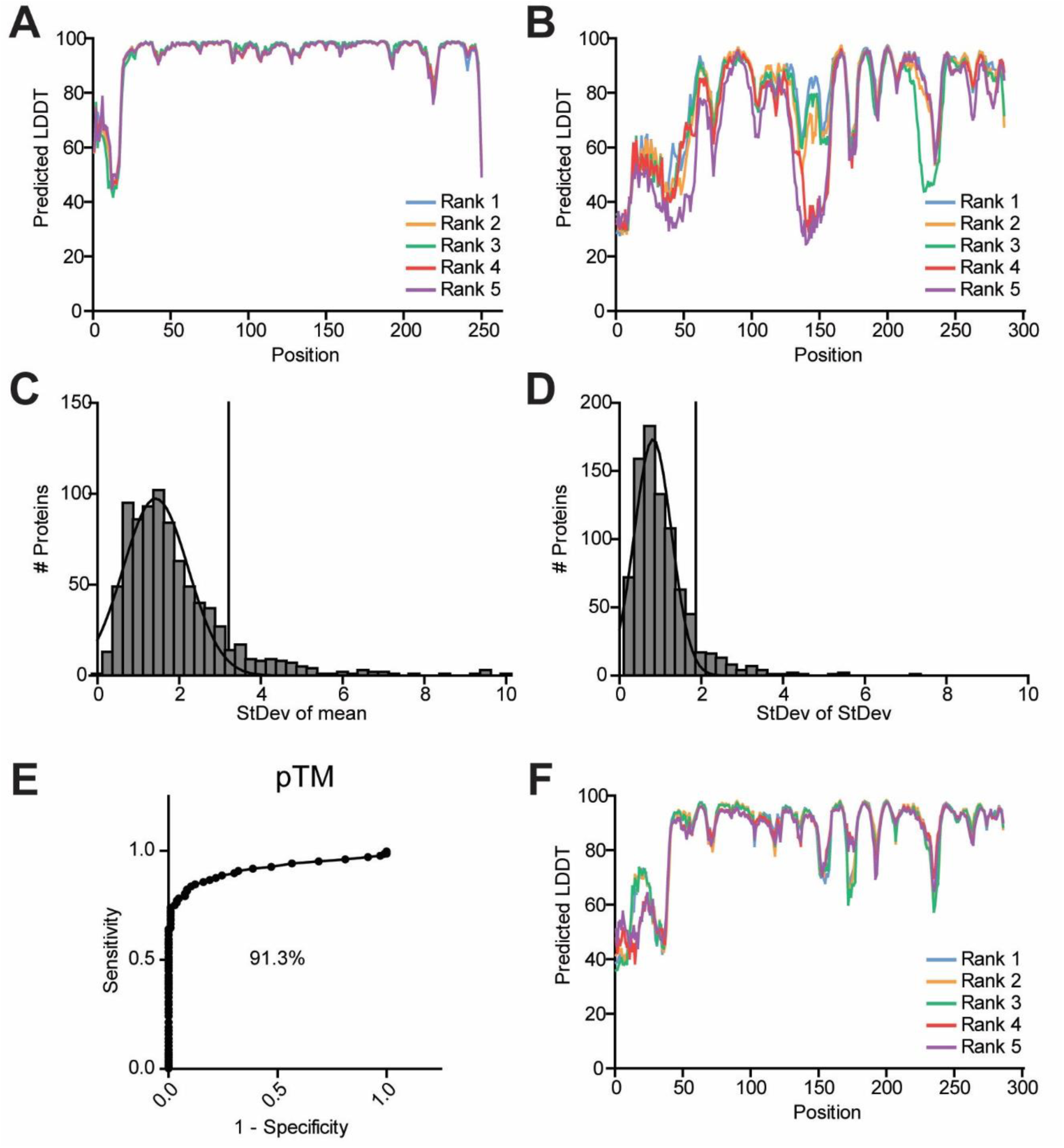
Developing thresholds for evaluating model quality. A. pLDDT plot of HCMV UL114 as an example of a protein where the pLDDT curves of the 5 models are consistent. B. pLDDT plot of HHV-7 U29 as an example of a protein where the pLDDT curves of the 5 models are inconsistent. C. Histogram of the standard deviation (StDev) of the mean of the pLDDT for each model for all LocalColabFold models. The threshold (black vertical bar) is set at the mean + 2x StDev, 3.2. D. Histogram of the StDev of the StDev of the pLDDT for each model for all LocalColabFold models. The threshold (black vertical bar) is set at the mean + 2x StDev, 1.9. E. ROC analysis of the pTM of all LocalColabFold models that passed the quality scores versus a negative control harboring scrambled amino acid sequences. The area under the curve is 91.3%. F. pLDDT curve of HHV-7 U29 as an example of a protein that passed the quality score thresholds after AlphaFold was used.

**Supplementary Fig. 2.**
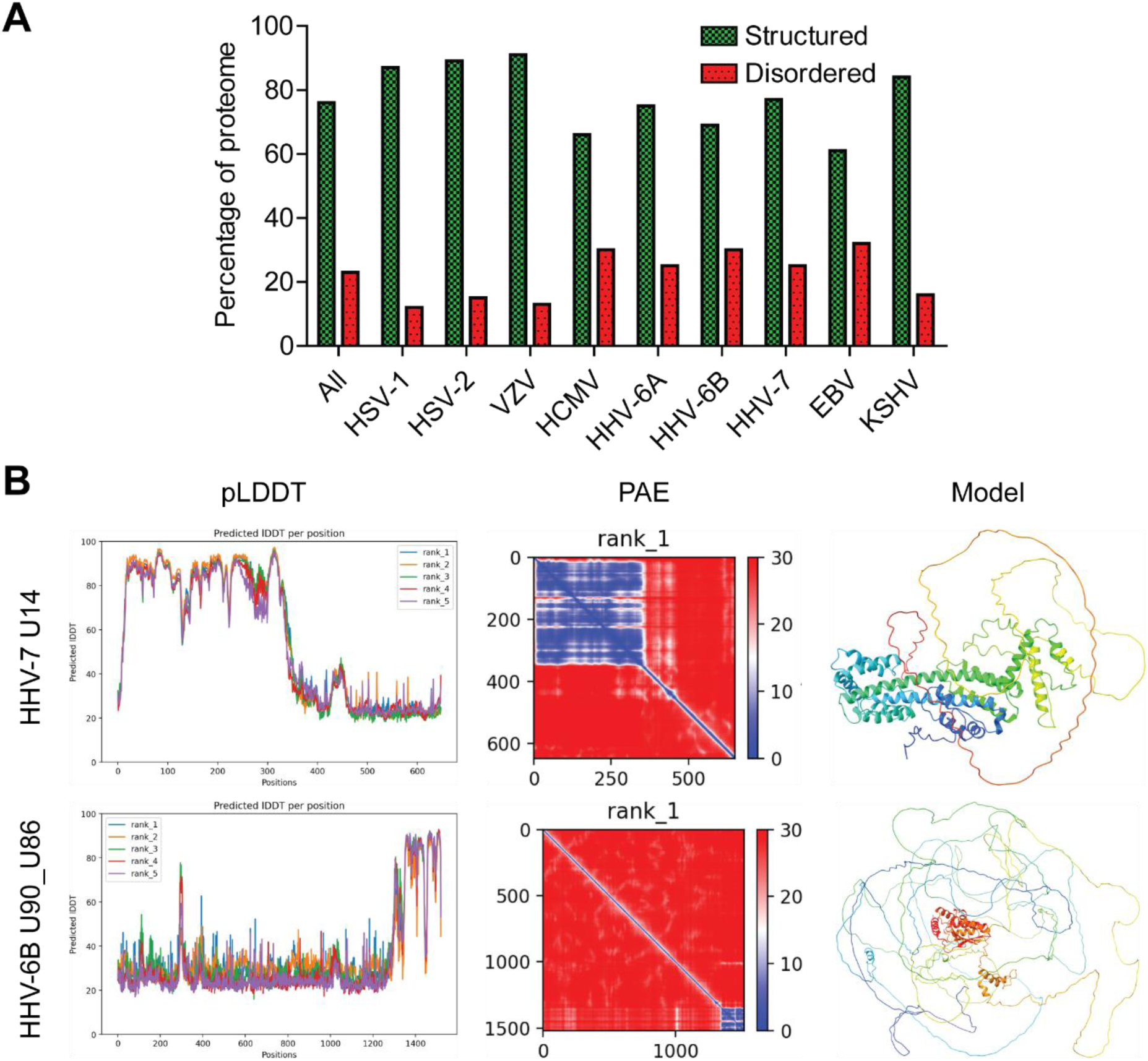
Herpesviruses express a large fraction of proteins that contain a disordered domain. A. The percentage of proteins that contain a structured or disordered domain are shown for each species as well as the consolidated proteome. B. HHV-7 U14 is an example of a protein with a structured N-terminus while HHV-6B U90_U86 has a structured C-terminus. The pLDDT and PAE plots are also shown to illustrate how the structured and disordered regions appear. The predicted structure is shown as a rainbow colored ribbon diagram where blue is the N-terminus and red is the C-terminus.

**Supplementary Fig. 3.**
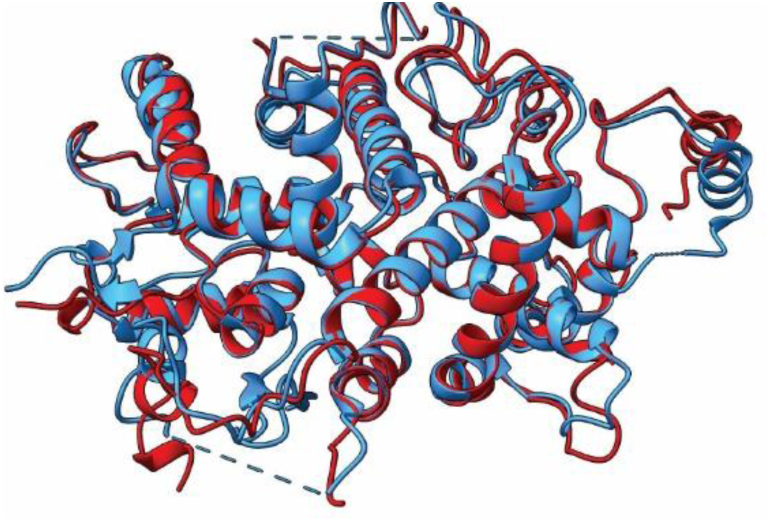
Alignment of predicted HCMV UL77 structure to the experimental structure that was deposited post-AlphaFold training. The deposited structure for HCMV UL77 (PDB 7NXP^19^) (blue) was aligned with the AlphaFold prediction (red), and the comparison has an RMSD of 0.509 Å against the 310 Cα. Only the residues present in the deposited structure are shown in the predicted model. The corresponding UCSF ChimeraX session can be found at https://zenodo.org/records/13284140.

**Supplementary Fig. 4.**
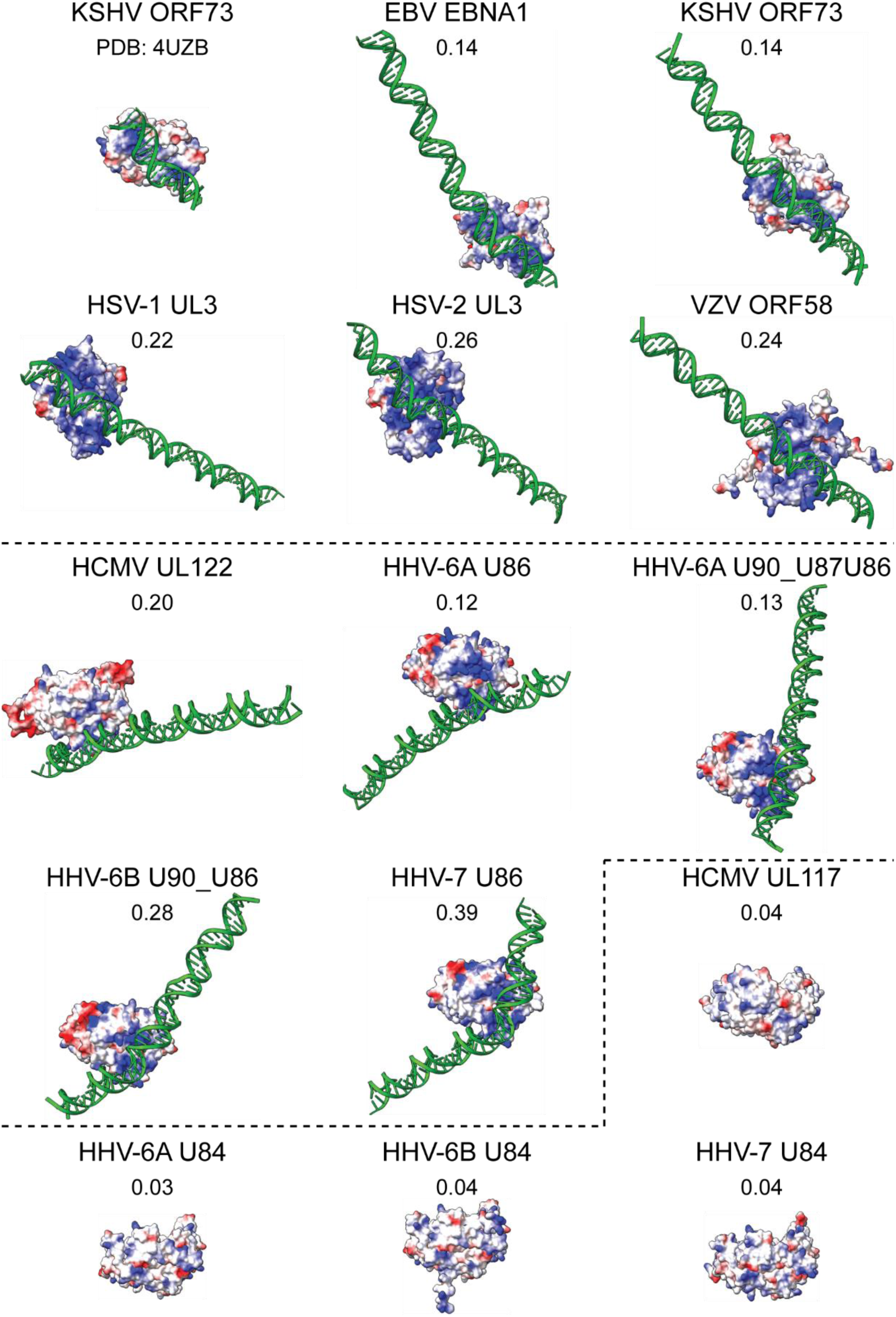
Predicted positive surface charge and DNA binding activity of the KSHV ORF73 and EBV EBNA1 cluster. The proteins in this cluster were structurally predicted as dimers of the conserved core with the DNA binding sequence of KSHV ORF73^80^ with AlphaFold 3. The crystal structure of KSHV ORF73 with DNA (PDB: 4UZB^29^) was included for comparison. The average pairwise ipTM between each protein and DNA strand is shown above the structure. The dashed line separates the different binding phenotypes: KSHV ORF73, HCMV UL122 (different orientation), and HCMV UL117 (no binding). Positive surface charge is displayed as blue, and negative charge is displayed as red. The corresponding UCSF ChimeraX session can be found at https://zenodo.org/records/13284140.

**Supplementary Fig. 5.**
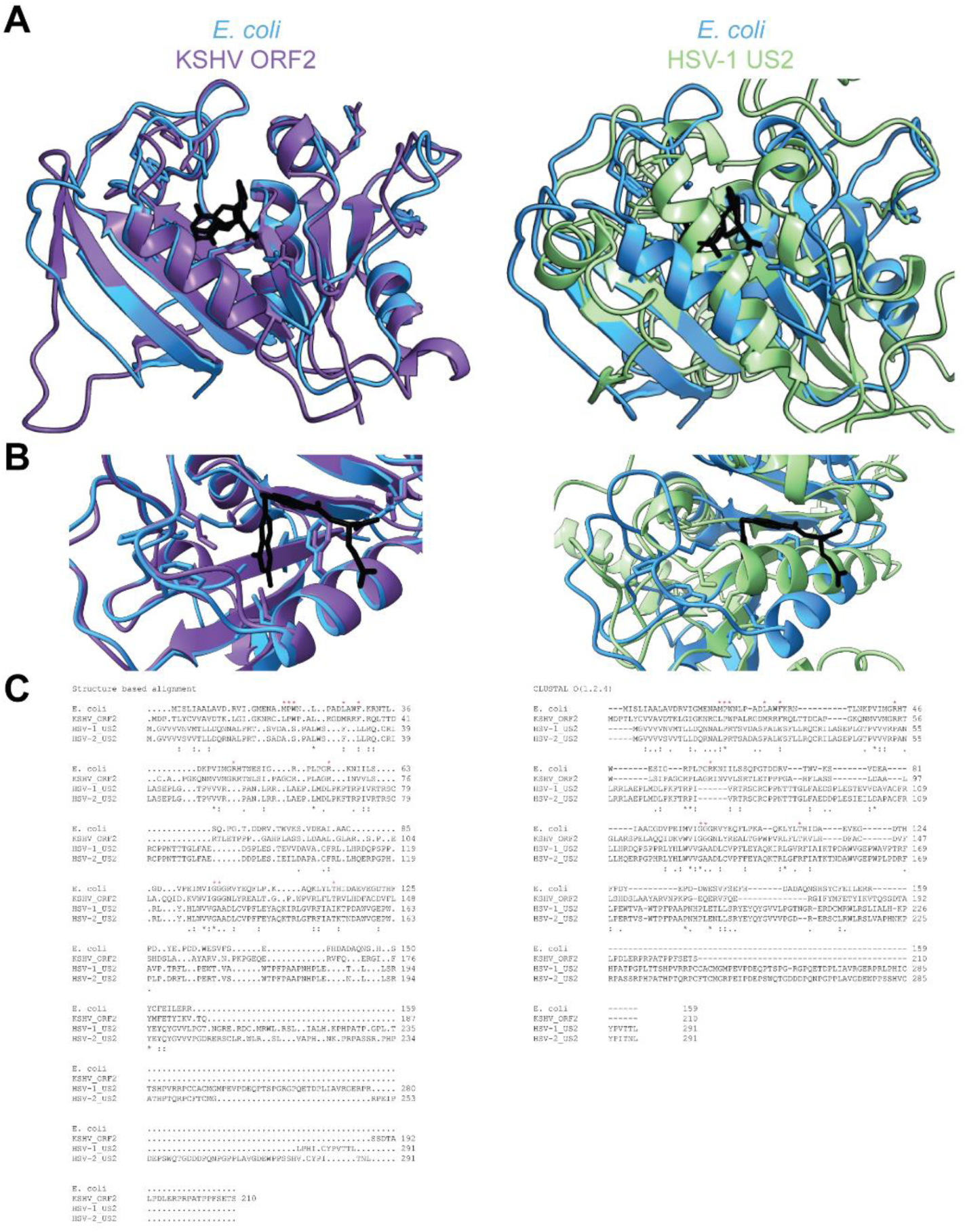
Alignment of *E. coli* DHFR to KSHV ORF2 and HSV-1 US2. A. The *E. coli* DHFR (PDB 1RX2^48^) (blue) was aligned to KSHV ORF2 (purple) and HSV-1 US2 (green), and the cofactor folic acid is shown in black. The atomic structure of the catalytic residues of the *E. coli* DHFR (M20, P21, W22, D27, F31, R44, R57, G95, G96, T113) and the potential catalytic residues that are in similar positions in ORF2 and US2 are shown. B. An enlarged view of the catalytic site is shown. C. The protein sequences were aligned based on the structure-based alignment or with Clustal Omega. The *E. coli* catalytic residues are marked with red asterisks. The potential catalytic residues in the herpesvirus proteins are from the structured based alignment. The corresponding UCSF ChimeraX session can be found at https://zenodo.org/records/13284140.

**Supplementary Fig. 6.**
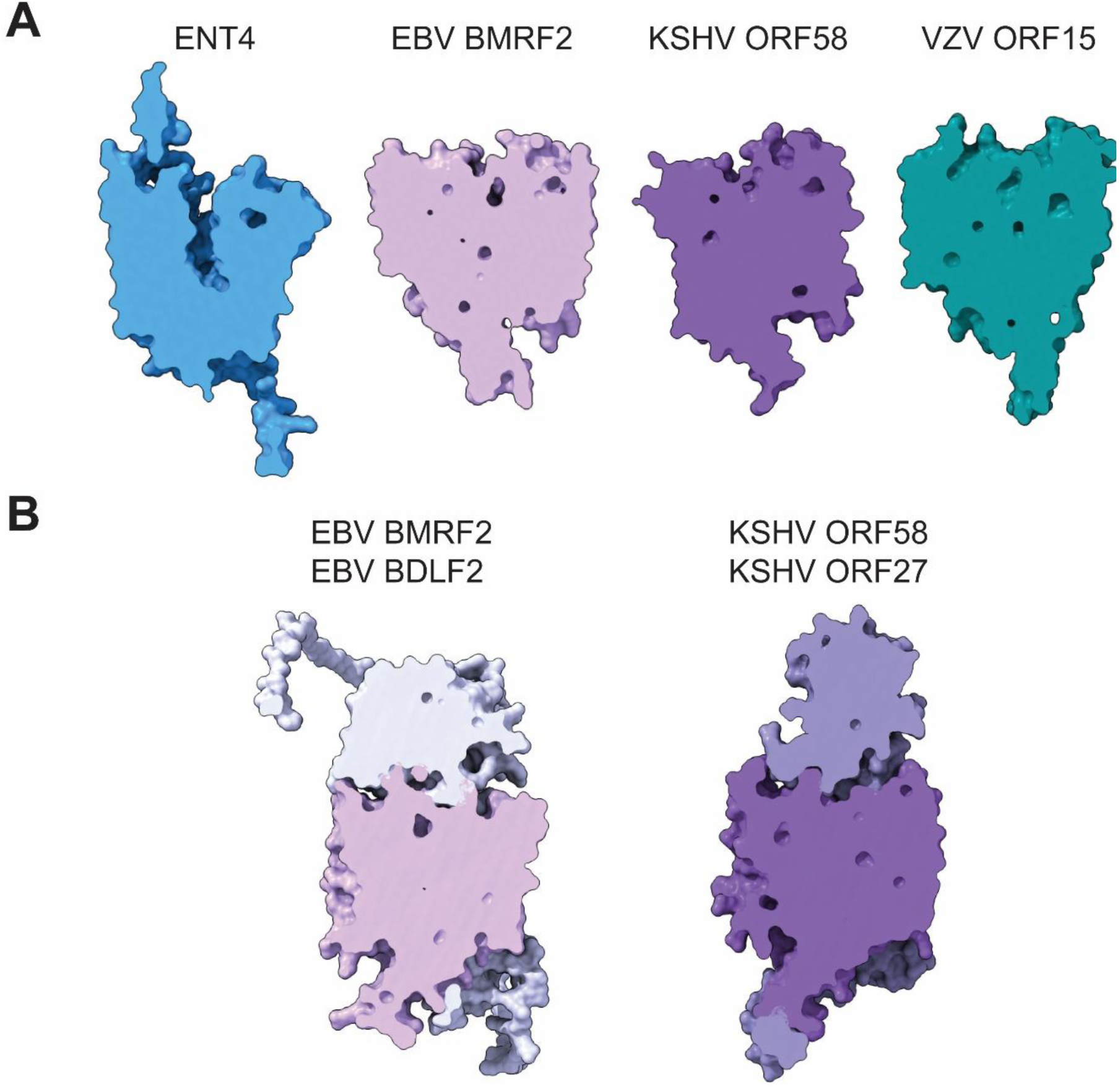
The herpesvirus ENTs are unlikely to have transporter activity. A. Cross-sections of surface plots of the monomer predictions of ENT4 (AF-Q8R139-F1-v4) and the structurally similar viral proteins. The white arrow points to the metabolite channel in ENT4. B. Structure predictions of the known EBV (BMRF2 pink and BDLF2 white) and KSHV (ORF58 purple and ORF27 light purple) heterodimers. The corresponding UCSF ChimeraX sessions can be found at https://zenodo.org/records/13284140.

## SUPPLEMENTARY FILES

**Supplementary Data 1. Consistency of pLDDT scores of AlphaFold models.** The quality scores for LocalColabFold with 3x recycles are in Sheet 1, AlphaFold are in Sheet 2, and LocalColabFold with 20x recycles are in Sheet 3. Sheet 4 lists all proteins and details whether they passed the quality scores. Sheet 5 lists the percentages of each protein sequence with a pLDDT of <0.50 (likely disordered) and >0.70 (likely structured). Sheet 6 lists the DeepTMHMM topology prediction of all proteins.

**Supplementary Data 2. Identified herpesvirus structural similarity clusters.** Sheet 1 lists the currently known homologous proteins between herpesvirus species and the Foldseek outputs. Sheet 2 shows the clustering of the herpesvirus proteins using the sequence alignment based algorithm MMseqs2, and Sheet 3 shows the clustering using the sequence HMM-HMM based algorithm HHblits. Sheet 4 lists the newly identified structural similarity groups based on DALI analysis with reference to the previously known homolog groups shown in Sheet 1. The identified function based on structural similarity to cellular structures is annotated. Sheet 5 is the same as Sheet 4, except that Foldseek outputs were used for clustering by structural similarity. Sheet 6 lists the structural clusters based on domain-level search instead of full-length protein search. The difference to the full length analysis is annotated. Sheet 7 shows the evaluation of the different Foldseek parameters. Specifically, it compares the different alignment algorithms, significance metrics, and significance thresholds. Sheet 8 compares the differences between analyzing full length herpesvirus proteins with Foldseek using the E-value versus the probability. The differences are highlighted. Sheet 9 compares the difference between analysing full-length proteins and proteins truncated based on the pLDDT score with Foldseek. The differences are highlighted. Sheet 10 lists all output protein pairs from HHblits and Foldseek along with the significance scores. A “-” denotes that the protein pair was not exported by the algorithm. For the significant pairs, they were classified as significant with only HHblits, with only Foldseek, or with both. Each protein pair has a score for protein A against B as well as B against A.

**Supplementary Data 3. Results of the domain-level structural similarity search.** Sheet 1 lists all pairwise similarities found using Foldseek for full-length similarity searches (columns A-B) and domain-level searches using structural snippets generated by the sliding window approach (columns C-D) for comparison. Pairs only identified in the domain-level search are listed again (columns E-F). Sheet 2 is a list of the sequence ranges of query proteins that match the sequence ranges of a target protein in the domain-level search. Sheet 3 lists all domain-level search results as well as a categorization of whether the hits can be categorized into three categories: "Internal Duplication" (the query protein matched itself), "Repetitive Acquisition" (the query protein matched a target sequence multiple times), or "Domain Addition" (the query protein has a domain that is not found in a related protein).

**Supplementary Data 4. Structural similarity search against cellular proteins.** Sheet 1 lists the Foldseek search results of the viral protein structure predictions against the PDB database structural similarity search. Sheet 2 lists the frequency of each word in the description of the hits from sheet 1 for each protein (column B). Viral keywords were subtracted to remove known functions and self-hits from experimental viral structures in the PDB (column C). Sheet 3 lists the Foldseek structural similarity search results of the viral protein structure predictions against the AlphaFold-Swiss-Prot database. Sheet 4 lists the frequency of each word in the description of the hits from sheet 3 for each protein (column B). Viral keywords were subtracted to remove known functions and self-hits from structures in the AlphaFold-Swiss-Prot database (column C). Sheet 5 lists which herpes proteins had a structurally similar herpes (column A) or cellular (column B) protein or is structurally unique (column C). It also lists viral clusters for which the whole cluster did not have any structurally similar proteins in the AlphaFold-Swiss-Prot database (column D). The PDB database was left out of the comparison for this sheet because the PDB also contains experimentally solved herpesvirus structures.

## References

1 Jumper, J., Evans, R., Pritzel, A., Green, T., Figurnov, M., Ronneberger, O., Tunyasuvunakool, K., Bates, R., Zidek, A., Potapenko, A., Bridgland, A., Meyer, C., Kohl, S. A. A., Ballard, A. J., Cowie, A., Romera-Paredes, B., Nikolov, S., Jain, R., Adler, J., Back, T., Petersen, S., Reiman, D., Clancy, E., Zielinski, M., Steinegger, M., Pacholska, M., Berghammer, T., Bodenstein, S., Silver, D., Vinyals, O., Senior, A. W., Kavukcuoglu, K., Kohli, P. & Hassabis, D. Highly accurate protein structure prediction with AlphaFold. Nature 596, 583–589, doi:10.1038/s41586-021-03819-2 (2021).

2 Baek, M., DiMaio, F., Anishchenko, I., Dauparas, J., Ovchinnikov, S., Lee, G. R., Wang, J., Cong, Q., Kinch, L. N., Schaeffer, R. D., Millan, C., Park, H., Adams, C., Glassman, C. R., DeGiovanni, A., Pereira, J. H., Rodrigues, A. V., van Dijk, A. A., Ebrecht, A. C., Opperman, D. J., Sagmeister, T., Buhlheller, C., Pavkov-Keller, T., Rathinaswamy, M. K., Dalwadi, U., Yip, C. K., Burke, J. E., Garcia, K. C., Grishin, N. V., Adams, P. D., Read, R. J. & Baker, D. Accurate prediction of protein structures and interactions using a three-track neural network. Science 373, 871–876, doi:10.1126/science.abj8754 (2021).

3 Akdel, M., Pires, D. E. V., Pardo, E. P., Janes, J., Zalevsky, A. O., Meszaros, B., Bryant, P., Good, L. L., Laskowski, R. A., Pozzati, G., Shenoy, A., Zhu, W., Kundrotas, P., Serra, V. R., Rodrigues, C. H. M., Dunham, A. S., Burke, D., Borkakoti, N., Velankar, S., Frost, A., Basquin, J., Lindorff-Larsen, K., Bateman, A., Kajava, A. V., Valencia, A., Ovchinnikov, S., Durairaj, J., Ascher, D. B., Thornton, J. M., Davey, N. E., Stein, A., Elofsson, A., Croll, T. I. & Beltrao, P. A structural biology community assessment of AlphaFold2 applications. Nat Struct Mol Biol 29, 1056–1067, doi:10.1038/s41594-022-00849-w (2022).

4 Zmasek, C. M., Knipe, D. M., Pellett, P. E. & Scheuermann, R. H. Classification of human Herpesviridae proteins using Domain-architecture Aware Inference of Orthologs (DAIO). Virology 529, 29–42, doi:10.1016/j.virol.2019.01.005 (2019).

5 van Kempen, M., Kim, S. S., Tumescheit, C., Mirdita, M., Lee, J., Gilchrist, C. L. M., Soding, J. & Steinegger, M. Fast and accurate protein structure search with Foldseek. Nat Biotechnol 42, 243–246, doi:10.1038/s41587-023-01773-0 (2024).

6 Abrescia, N. G., Bamford, D. H., Grimes, J. M. & Stuart, D. I. Structure unifies the viral universe. Annu Rev Biochem 81, 795–822, doi:10.1146/annurev-biochem-060910-095130 (2012).

7 Ng, W. M., Stelfox, A. J. & Bowden, T. A. Unraveling virus relationships by structure-based phylogenetic classification. Virus Evol 6, veaa003, doi:10.1093/ve/veaa003 (2020).

8 Oliver, M. R., Toon, K., Lewis, C. B., Devlin, S., Gifford, R. J. & Grove, J. Structures of the Hepaci-, Pegi-, and Pestiviruses envelope proteins suggest a novel membrane fusion mechanism. PLoS Biol 21, e3002174, doi:10.1371/journal.pbio.3002174 (2023).

9 Evseev, P., Gutnik, D., Shneider, M. & Miroshnikov, K. Use of an Integrated Approach Involving AlphaFold Predictions for the Evolutionary Taxonomy of Duplodnaviria Viruses. Biomolecules 13, doi:10.3390/biom13010110 (2023).

10 Mutz, P., Resch, W., Faure, G., Senkevich, T. G., Koonin, E. V. & Moss, B. Exaptation of Inactivated Host Enzymes for Structural Roles in Orthopoxviruses and Novel Folds of Virus Proteins Revealed by Protein Structure Modeling. mBio 14, e0040823, doi:10.1128/mbio.00408-23 (2023).

11 Mirdita, M., Schutze, K., Moriwaki, Y., Heo, L., Ovchinnikov, S. & Steinegger, M. ColabFold: making protein folding accessible to all. Nat Methods 19, 679–682, doi:10.1038/s41592-022-01488-1 (2022).

12 Terwilliger, T. C., Liebschner, D., Croll, T. I., Williams, C. J., McCoy, A. J., Poon, B. K., Afonine, P. V., Oeffner, R. D., Richardson, J. S., Read, R. J. & Adams, P. D. AlphaFold predictions are valuable hypotheses and accelerate but do not replace experimental structure determination. Nat Methods 21, 110–116, doi:10.1038/s41592-023-02087-4 (2024).

13 Necci, M., Piovesan, D., Predictors, C., DisProt, C. & Tosatto, S. C. E. Critical assessment of protein intrinsic disorder prediction. Nat Methods 18, 472–481, doi:10.1038/s41592-021-01117-3 (2021).

14 Piovesan, D., Monzon, A. M. & Tosatto, S. C. E. Intrinsic protein disorder and conditional folding in AlphaFoldDB. Protein Sci 31, e4466, doi:10.1002/pro.4466 (2022).

15 Piovesan, D., Del Conte, A., Clementel, D., Monzon, A. M., Bevilacqua, M., Aspromonte, M. C., Iserte, J. A., Orti, F. E., Marino-Buslje, C. & Tosatto, S. C. E. MobiDB: 10 years of intrinsically disordered proteins. Nucleic Acids Res 51, D438–D444, doi:10.1093/nar/gkac1065 (2023).

16 Caragliano, E., Bonazza, S., Frascaroli, G., Tang, J., Soh, T. K., Grunewald, K., Bosse, J. B. & Brune, W. Human cytomegalovirus forms phase-separated compartments at viral genomes to facilitate viral replication. Cell Rep 38, 110469, doi:10.1016/j.celrep.2022.110469 (2022).

17 Dunker, A. K., Obradovic, Z., Romero, P., Garner, E. C. & Brown, C. J. Intrinsic protein disorder in complete genomes. Genome Inform Ser Workshop Genome Inform 11, 161–171 (2000).

18 Oldfield, C. J., Cheng, Y., Cortese, M. S., Brown, C. J., Uversky, V. N. & Dunker, A. K. Comparing and combining predictors of mostly disordered proteins. Biochemistry 44, 1989–2000, doi:10.1021/bi047993o (2005).

19 Naniima, P., Naimo, E., Koch, S., Curth, U., Alkharsah, K. R., Stroh, L. J., Binz, A., Beneke, J. M., Vollmer, B., Boning, H., Borst, E. M., Desai, P., Bohne, J., Messerle, M., Bauerfeind, R., Legrand, P., Sodeik, B., Schulz, T. F. & Krey, T. Assembly of infectious Kaposi’s sarcoma-associated herpesvirus progeny requires formation of a pORF19 pentamer. PLoS Biol 19, e3001423, doi:10.1371/journal.pbio.3001423 (2021).

20 The UniProt, C. UniProt: the universal protein knowledgebase. Nucleic Acids Res 45, D158–D169, doi:10.1093/nar/gkw1099 (2017).

21 in *Human Herpesviruses: Biology, Therapy, and Immunoprophylaxis* (eds A. Arvin et al.) (2007).

22 Holm, L. & Sander, C. Dali: a network tool for protein structure comparison. Trends Biochem Sci 20, 478–480, doi:10.1016/s0968-0004(00)89105-7 (1995).

23 Bermek, O. & Williams, R. S. The three-component helicase/primase complex of herpes simplex virus-1. Open Biol 11, 210011, doi:10.1098/rsob.210011 (2021).

24 Kazlauskas, D. & Venclovas, C. Herpesviral helicase-primase subunit UL8 is inactivated B-family polymerase. Bioinformatics 30, 2093–2097, doi:10.1093/bioinformatics/btu204 (2014).

25 Weisshart, K., Kuo, A. A., Hwang, C. B., Kumura, K. & Coen, D. M. Structural and functional organization of herpes simplex virus DNA polymerase investigated by limited proteolysis. J Biol Chem 269, 22788–22796 (1994).

26 Jahedi, S., Markovitz, N. S., Filatov, F. & Roizman, B. Colocalization of the herpes simplex virus 1 UL4 protein with infected cell protein 22 in small, dense nuclear structures formed prior to onset of DNA synthesis. J Virol 73, 5132–5138, doi:10.1128/JVI.73.6.5132-5136.1999 (1999).

27 Bochkarev, A., Barwell, J. A., Pfuetzner, R. A., Bochkareva, E., Frappier, L. & Edwards, A. M. Crystal structure of the DNA-binding domain of the Epstein-Barr virus origin-binding protein, EBNA1, bound to DNA. Cell 84, 791–800, doi:10.1016/s0092-8674(00)81056-9 (1996).

28 Stevenson, J. P. Correspondence: Fish diseases research. Vet Rec 95, 325–326, doi:10.1136/vr.95.14.325 (1974).

29 Hellert, J., Weidner-Glunde, M., Krausze, J., Lunsdorf, H., Ritter, C., Schulz, T. F. & Luhrs, T. The 3D structure of Kaposi sarcoma herpesvirus LANA C-terminal domain bound to DNA. Proc Natl Acad Sci U S A 112, 6694–6699, doi:10.1073/pnas.1421804112 (2015).

30 Ball, C. B., Li, M., Parida, M., Hu, Q., Ince, D., Collins, G. S., Meier, J. L. & Price, D. H. Human Cytomegalovirus IE2 Both Activates and Represses Initiation and Modulates Elongation in a Context-Dependent Manner. mBio 13, e0033722, doi:10.1128/mbio.00337-22 (2022).

31 Gravel, A., Tomoiu, A., Cloutier, N., Gosselin, J. & Flamand, L. Characterization of the immediate-early 2 protein of human herpesvirus 6, a promiscuous transcriptional activator. Virology 308, 340–353, doi:10.1016/s0042-6822(03)00007-2 (2003).

32 Szymanska-de Wijs, K., Dezeljin, M., Bogdanow, B. & Messerle, M. Viral determinants influencing intra-and intercellular communication in cytomegalovirus infection. Curr Opin Virol 60, 101328, doi:10.1016/j.coviro.2023.101328 (2023).

33 Bogdanow, B., Gruska, I., Muhlberg, L., Protze, J., Hohensee, S., Vetter, B., Bosse, J. B., Lehmann, M., Sadeghi, M., Wiebusch, L. & Liu, F. Spatially resolved protein map of intact human cytomegalovirus virions. Nat Microbiol 8, 1732–1747, doi:10.1038/s41564-023-01433-8 (2023).

34 Nabiee, R., Syed, B., Ramirez Castano, J., Lalani, R. & Totonchy, J. E. An Update of the Virion Proteome of Kaposi Sarcoma-Associated Herpesvirus. Viruses 12, doi:10.3390/v12121382 (2020).

35 Davison, A. J., Akter, P., Cunningham, C., Dolan, A., Addison, C., Dargan, D. J., Hassan-Walker, A. F., Emery, V. C., Griffiths, P. D. & Wilkinson, G. W. G. Homology between the human cytomegalovirus RL11 gene family and human adenovirus E3 genes. J Gen Virol 84, 657–663, doi:10.1099/vir.0.18856-0 (2003).

36 Dunn, W., Chou, C., Li, H., Hai, R., Patterson, D., Stolc, V., Zhu, H. & Liu, F. Functional profiling of a human cytomegalovirus genome. Proc Natl Acad Sci U S A 100, 14223–14228, doi:10.1073/pnas.2334032100 (2003).

37 Osanyinlusi, S. A., Zischke, J., Jacobs, R., Weissinger, E. M., Schulz, T. F. & Kay-Fedorov, P. C. Human Cytomegalovirus pUL11, a CD45 Ligand, Disrupts CD4 T Cell Control of Viral Spread in Epithelial Cells. mBio 13, e0294622, doi:10.1128/mbio.02946-22 (2022).

38 Chee, M. S., Bankier, A. T., Beck, S., Bohni, R., Brown, C. M., Cerny, R., Horsnell, T., Hutchison, C. A., 3rd, Kouzarides, T., Martignetti, J. A. & et al. Analysis of the protein-coding content of the sequence of human cytomegalovirus strain AD169. Curr Top Microbiol Immunol 154, 125–169, doi:10.1007/978-3-642-74980-3_6 (1990).

39 Menard, C., Wagner, M., Ruzsics, Z., Holak, K., Brune, W., Campbell, A. E. & Koszinowski, U. H. Role of murine cytomegalovirus US22 gene family members in replication in macrophages. J Virol 77, 5557–5570, doi:10.1128/jvi.77.10.5557-5570.2003 (2003).

40 Buisson, M., Geoui, T., Flot, D., Tarbouriech, N., Ressing, M. E., Wiertz, E. J. & Burmeister, W. P. A bridge crosses the active-site canyon of the Epstein-Barr virus nuclease with DNase and RNase activities. J Mol Biol 391, 717–728, doi:10.1016/j.jmb.2009.06.034 (2009).

41 Dahlroth, S. L., Gurmu, D., Schmitzberger, F., Engman, H., Haas, J., Erlandsen, H. & Nordlund, P. Crystal structure of the shutoff and exonuclease protein from the oncogenic Kaposi’s sarcoma-associated herpesvirus. FEBS J 276, 6636–6645, doi:10.1111/j.1742-4658.2009.07374.x (2009).

42 Lee, H., Patschull, A. O. M., Bagneris, C., Ryan, H., Sanderson, C. M., Ebrahimi, B., Nobeli, I. & Barrett, T. E. KSHV SOX mediated host shutoff: the molecular mechanism underlying mRNA transcript processing. Nucleic Acids Res 45, 4756–4767, doi:10.1093/nar/gkw1340 (2017).

43 Maninger, S., Bosse, J. B., Lemnitzer, F., Pogoda, M., Mohr, C. A., von Einem, J., Walther, P., Koszinowski, U. H. & Ruzsics, Z. M94 is essential for the secondary envelopment of murine cytomegalovirus. J Virol 85, 9254–9267, doi:10.1128/JVI.00443-11 (2011).

44 Volpato, J. P., Yachnin, B. J., Blanchet, J., Guerrero, V., Poulin, L., Fossati, E., Berghuis, A. M. & Pelletier, J. N. Multiple conformers in active site of human dihydrofolate reductase F31R/Q35E double mutant suggest structural basis for methotrexate resistance. J Biol Chem 284, 20079–20089, doi:10.1074/jbc.M109.018010 (2009).

45 Sarid, R., Flore, O., Bohenzky, R. A., Chang, Y. & Moore, P. S. Transcription mapping of the Kaposi’s sarcoma-associated herpesvirus (human herpesvirus 8) genome in a body cavity-based lymphoma cell line (BC-1). J Virol 72, 1005–1012, doi:10.1128/JVI.72.2.1005-1012.1998 (1998).

46 Nicholas, J., Zong, J. C., Alcendor, D. J., Ciufo, D. M., Poole, L. J., Sarisky, R. T., Chiou, C. J., Zhang, X., Wan, X., Guo, H. G., Reitz, M. S. & Hayward, G. S. Novel organizational features, captured cellular genes, and strain variability within the genome of KSHV/HHV8. J Natl Cancer Inst Monogr, 79–88, doi:10.1093/oxfordjournals.jncimonographs.a024179 (1998).

47 Howell, E. E. Searching sequence space: two different approaches to dihydrofolate reductase catalysis. Chembiochem 6, 590–600, doi:10.1002/cbic.200400237 (2005).

48 Sawaya, M. R. & Kraut, J. Loop and subdomain movements in the mechanism of Escherichia coli dihydrofolate reductase: crystallographic evidence. Biochemistry 36, 586–603, doi:10.1021/bi962337c (1997).

49 Barnes, K., Dobrzynski, H., Foppolo, S., Beal, P. R., Ismat, F., Scullion, E. R., Sun, L., Tellez, J., Ritzel, M. W., Claycomb, W. C., Cass, C. E., Young, J. D., Billeter-Clark, R., Boyett, M. R. & Baldwin, S. A. Distribution and functional characterization of equilibrative nucleoside transporter-4, a novel cardiac adenosine transporter activated at acidic pH. Circ Res 99, 510–519, doi:10.1161/01.RES.0000238359.18495.42 (2006).

50 Loesing, J. B., Di Fiore, S., Ritter, K., Fischer, R. & Kleines, M. Epstein-Barr virus BDLF2-BMRF2 complex affects cellular morphology. J Gen Virol 90, 1440–1449, doi:10.1099/vir.0.009571-0 (2009).

51 Walston, J. J., Hayman, I. R., Gore, M., Ferguson, M., Temple, R. M., Liao, J., Alam, S., Meyers, C., Tugizov, S. M., Hutt-Fletcher, L. & Sample, C. E. The Epstein-Barr Virus Glycoprotein BDLF2 Is Essential for Efficient Viral Spread in Stratified Epithelium. J Virol 97, e0152822, doi:10.1128/jvi.01528-22 (2023).

52 Klupp, B. G., Altenschmidt, J., Granzow, H., Fuchs, W. & Mettenleiter, T. C. Identification and characterization of the pseudorabies virus UL43 protein. Virology 334, 224–233, doi:10.1016/j.virol.2005.01.032 (2005).

53 El Kasmi, I. & Lippe, R. Herpes simplex virus 1 gN partners with gM to modulate the viral fusion machinery. J Virol 89, 2313–2323, doi:10.1128/JVI.03041-14 (2015).

54 Wright, N. J. & Lee, S. Y. Structures of human ENT1 in complex with adenosine reuptake inhibitors. Nat Struct Mol Biol 26, 599–606, doi:10.1038/s41594-019-0245-7 (2019).

55 Massa Lopez, D., Thelen, M., Stahl, F., Thiel, C., Linhorst, A., Sylvester, M., Hermanns-Borgmeyer, I., Lullmann-Rauch, R., Eskild, W., Saftig, P. & Damme, M. The lysosomal transporter MFSD1 is essential for liver homeostasis and critically depends on its accessory subunit GLMP. Elife 8, doi:10.7554/eLife.50025 (2019).

56 Nomburg, J., Doherty, E. E., Price, N., Bellieny-Rabelo, D., Zhu, Y. K. & Doudna, J. A. Birth of protein folds and functions in the virome. Nature 633, 710–717, doi:10.1038/s41586-024-07809-y (2024).

57 Davison, A. J. & Stow, N. D. New genes from old: redeployment of dUTPase by herpesviruses. J Virol 79, 12880–12892, doi:10.1128/JVI.79.20.12880-12892.2005 (2005).

58 Tarbouriech, N., Buisson, M., Seigneurin, J. M., Cusack, S. & Burmeister, W. P. The monomeric dUTPase from Epstein-Barr virus mimics trimeric dUTPases. Structure 13, 1299–1310, doi:10.1016/j.str.2005.06.009 (2005).

59 Williams, M. V., Cox, B. & Ariza, M. E. Herpesviruses dUTPases: A New Family of Pathogen-Associated Molecular Pattern (PAMP) Proteins with Implications for Human Disease. Pathogens 6, doi:10.3390/pathogens6010002 (2016).

60 Eberhage, J., Bresch, I. P., Ramani, R., Viohl, N., Buchta, T., Rehfeld, C. L., Hinse, P., Reubold, T. F., Brinkmann, M. M. & Eschenburg, S. Crystal structure of the tegument protein UL82 (pp71) from human cytomegalovirus. Protein Sci 33, e4915, doi:10.1002/pro.4915 (2024).

61 Zhao, Z. S., Granucci, F., Yeh, L., Schaffer, P. A. & Cantor, H. Molecular mimicry by herpes simplex virus-type 1: autoimmune disease after viral infection. Science 279, 1344–1347, doi:10.1126/science.279.5355.1344 (1998).

62 Soldan, S. S. & Lieberman, P. M. Epstein-Barr virus and multiple sclerosis. Nat Rev Microbiol 21, 51–64, doi:10.1038/s41579-022-00770-5 (2023).

63 Begum, S., Aiman, S., Ahmad, S., Samad, A., Almehmadi, M., Allahyani, M., Aljuaid, A., Afridi, S. G. & Khan, A. Molecular Mimicry Analyses Unveiled the Human Herpes Simplex and Poxvirus Epitopes as Possible Candidates to Incite Autoimmunity. Pathogens 11, doi:10.3390/pathogens11111362 (2022).

64 Lunemann, J. D., Jelcic, I., Roberts, S., Lutterotti, A., Tackenberg, B., Martin, R. & Munz, C. EBNA1-specific T cells from patients with multiple sclerosis cross react with myelin antigens and co-produce IFN-gamma and IL-2. J Exp Med 205, 1763–1773, doi:10.1084/jem.20072397 (2008).

65 Heldwein, E. E., Lou, H., Bender, F. C., Cohen, G. H., Eisenberg, R. J. & Harrison, S. C. Crystal structure of glycoprotein B from herpes simplex virus 1. Science 313, 217–220, doi:10.1126/science.1126548 (2006).

66 Hekkelman, M. L., de Vries, I., Joosten, R. P. & Perrakis, A. AlphaFill: enriching AlphaFold models with ligands and cofactors. Nat Methods 20, 205–213, doi:10.1038/s41592-022-01685-y (2023).

67 Pang, M., He, W., Lu, X., She, Y., Xie, L., Kong, R. & Chang, S. CoDock-Ligand: combined template-based docking and CNN-based scoring in ligand binding prediction. BMC Bioinformatics 24, 444, doi:10.1186/s12859-023-05571-y (2023).

68 Baek, M., McHugh, R., Anishchenko, I., Jiang, H., Baker, D. & DiMaio, F. Accurate prediction of protein-nucleic acid complexes using RoseTTAFoldNA. Nat Methods 21, 117–121, doi:10.1038/s41592-023-02086-5 (2024).

69 Krishna, R., Wang, J., Ahern, W., Sturmfels, P., Venkatesh, P., Kalvet, I., Lee, G. R., Morey-Burrows, F. S., Anishchenko, I., Humphreys, I. R., McHugh, R., Vafeados, D., Li, X., Sutherland, G. A., Hitchcock, A., Hunter, C. N., Kang, A., Brackenbrough, E., Bera, A. K., Baek, M., DiMaio, F. & Baker, D. Generalized biomolecular modeling and design with RoseTTAFold All-Atom. Science 384, eadl2528, doi:10.1126/science.adl2528 (2024).

70 Abramson, J., Adler, J., Dunger, J., Evans, R., Green, T., Pritzel, A., Ronneberger, O., Willmore, L., Ballard, A. J., Bambrick, J., Bodenstein, S. W., Evans, D. A., Hung, C. C., O’Neill, M., Reiman, D., Tunyasuvunakool, K., Wu, Z., Zemgulyte, A., Arvaniti, E., Beattie, C., Bertolli, O., Bridgland, A., Cherepanov, A., Congreve, M., Cowen-Rivers, A. I., Cowie, A., Figurnov, M., Fuchs, F. B., Gladman, H., Jain, R., Khan, Y. A., Low, C. M. R., Perlin, K., Potapenko, A., Savy, P., Singh, S., Stecula, A., Thillaisundaram, A., Tong, C., Yakneen, S., Zhong, E. D., Zielinski, M., Zidek, A., Bapst, V., Kohli, P., Jaderberg, M., Hassabis, D. & Jumper, J. M. Accurate structure prediction of biomolecular interactions with AlphaFold 3. Nature 630, 493–500, doi:10.1038/s41586-024-07487-w (2024).

71 Whisnant, A. W., Jurges, C. S., Hennig, T., Wyler, E., Prusty, B., Rutkowski, A. J., L’Hernault, A., Djakovic, L., Gobel, M., Doring, K., Menegatti, J., Antrobus, R., Matheson, N. J., Kunzig, F. W. H., Mastrobuoni, G., Bielow, C., Kempa, S., Liang, C., Dandekar, T., Zimmer, R., Landthaler, M., Grasser, F., Lehner, P. J., Friedel, C. C., Erhard, F. & Dolken, L. Integrative functional genomics decodes herpes simplex virus 1. Nat Commun 11, 2038, doi:10.1038/s41467-020-15992-5 (2020).

72 Stern-Ginossar, N., Weisburd, B., Michalski, A., Le, V. T., Hein, M. Y., Huang, S. X., Ma, M., Shen, B., Qian, S. B., Hengel, H., Mann, M., Ingolia, N. T. & Weissman, J. S. Decoding human cytomegalovirus. Science 338, 1088–1093, doi:10.1126/science.1227919 (2012).

73 Bencun, M., Klinke, O., Hotz-Wagenblatt, A., Klaus, S., Tsai, M. H., Poirey, R. & Delecluse, H. J. Translational profiling of B cells infected with the Epstein-Barr virus reveals 5’ leader ribosome recruitment through upstream open reading frames. Nucleic Acids Res 46, 2802–2819, doi:10.1093/nar/gky129 (2018).

74 Arias, C., Weisburd, B., Stern-Ginossar, N., Mercier, A., Madrid, A. S., Bellare, P., Holdorf, M., Weissman, J. S. & Ganem, D. KSHV 2.0: a comprehensive annotation of the Kaposi’s sarcoma-associated herpesvirus genome using next-generation sequencing reveals novel genomic and functional features. PLoS Pathog 10, e1003847, doi:10.1371/journal.ppat.1003847 (2014).

75 Shekhar, R., O’Grady, T., Keil, N., Feswick, A., Amador, D. A. M., Tibbetts, S. A., Flemington, E. K. & Renne, R. High-density resolution of the Kaposi’s sarcoma associated herpesvirus transcriptome identifies novel transcript isoforms generated by long-range transcription and alternative splicing. Nucleic Acids Res 52, 7720–7739, doi:10.1093/nar/gkae540 (2024).

76 Steinegger, M. & Soding, J. MMseqs2 enables sensitive protein sequence searching for the analysis of massive data sets. Nat Biotechnol 35, 1026–1028, doi:10.1038/nbt.3988 (2017).

77 Remmert, M., Biegert, A., Hauser, A. & Soding, J. HHblits: lightning-fast iterative protein sequence searching by HMM-HMM alignment. Nat Methods 9, 173–175, doi:10.1038/nmeth.1818 (2011).

78 Hallgren, J., Tsirigos, K. D., Pedersen, M. D., Armenteros, J. J. A., Marcatili, P., Nielsen, H., Krogh, A. & Winther, O. DeepTMHMM predicts alpha and beta transmembrane proteins using deep neural networks. bioRxiv, doi:10.1101/2022.04.08.487609 (2022).

79 Holm, L. Using Dali for Protein Structure Comparison. Methods Mol Biol 2112, 29–42, doi:10.1007/978-1-0716-0270-6_3 (2020).

80 Hu, J., Garber, A. C. & Renne, R. The latency-associated nuclear antigen of Kaposi’s sarcoma-associated herpesvirus supports latent DNA replication in dividing cells. J Virol 76, 11677–11687, doi:10.1128/jvi.76.22.11677-11687.2002 (2002).

